# High coverage whole genome sequencing of the expanded 1000 Genomes Project cohort including 602 trios

**DOI:** 10.1101/2021.02.06.430068

**Authors:** Marta Byrska-Bishop, Uday S. Evani, Xuefang Zhao, Anna O. Basile, Haley J. Abel, Allison A. Regier, André Corvelo, Wayne E. Clarke, Rajeeva Musunuri, Kshithija Nagulapalli, Susan Fairley, Alexi Runnels, Lara Winterkorn, Ernesto Lowy, The Human Genome Structural Variation Consortium, Paul Flicek, Soren Germer, Harrison Brand, Ira M. Hall, Michael E. Talkowski, Giuseppe Narzisi, Michael C. Zody

## Abstract

The 1000 Genomes Project (1kGP) is the largest fully open resource of whole genome sequencing (WGS) data consented for public distribution of raw sequence data without access or use restrictions. The final release of the 1kGP included 2,504 unrelated samples from 26 populations and was based primarily on low coverage WGS. Here, we present a new, *high coverage* 3,202-sample WGS 1kGP resource, sequenced to a targeted depth of 30X using the Illumina NovaSeq 6000 system, which now includes 602 complete trios. We performed SNV/INDEL calling against the GRCh38 reference using GATK’s HaplotypeCaller, and generated a comprehensive set of SVs by integrating multiple analytic methods through a sophisticated machine learning model. We make all the data generated as part of this project publicly available and we envision it to become the new de facto public resource for the worldwide genomics and genetics community.

## INTRODUCTION

The 1000 Genomes Project (1kGP) was the first large-scale whole genome sequencing (WGS) effort to deliver a catalog of human genetic variation (Sudmant et al., 2015; The 1000 Genomes Project Consortium, 2010, 2012, 2015). The project sampled participants from 26 populations across five continental regions of the world. Spanning seven years of data generation and analysis, it culminated in 2015 with a publication of the final, phase 3, variant call set (Sudmant et al., 2015; The 1000 Genomes Project Consortium, 2015) consisting of 2,504 unrelated samples, a subset of which is from the HapMap collection (The International HapMap 3 Consortium, 2010). The set of 2,504 samples was selected with the goal of maximizing the discovery of single nucleotide variants (SNVs) at minor allele frequencies (MAF) of 1% or higher in diverse populations, hence related samples were not included. The phase 3 call set was generated based on the combination of low coverage WGS (mean depth 7.4X), high-coverage whole exome sequencing (WES, mean depth 65.7X), and microarray genotyping data. It included 84.7 million SNVs, and 3.6 million short insertions and deletions (INDELs), as well as a separate set of 68,818 structural variants (SVs; alterations ≥ 50 bp). The 1kGP resources have been collectively cited over 16,000 times to date and have been utilized for foundational applications such as genotype imputation, eQTL mapping, variant pathogenicity prioritization, population history, and evolutionary genetics studies (Almeida et al., 2014; Hara et al., 2014; Horikoshi et al., 2015; Huang et al., 2015; Khurana et al., 2013; Kircher et al., 2014; Lappalainen et al., 2013; Nikpay et al., 2015; Ritchie et al., 2014; Zheng-Bradley and Flicek, 2017). While the phase 3 dataset captured the vast majority of common variants (MAF > 1%) in the population (> 99% of SNVs and > 85 % INDELs) (The 1000 Genomes Project Consortium, 2015), detection of rare variants (MAF ≤ 1%) was limited due to low sequencing coverage outside of the coding regions of the genome.

Here, we present high coverage WGS and comprehensive analyses of the original 2,504 1kGP samples, as well as of 698 additional related samples. These related samples were not included as part of the phase 3 call set, but now provide complete WGS on 602 trios in the 1kGP cohort. A small subset of these pedigrees have been sequenced previously as part of various efforts, such as Platinum Genomes (Eberle et al., 2017), Complete Genomics (The 1000 Genomes Project Consortium, 2015), and the Human Genome Structural Variation Consortium (HGSVC), which generated long-read WGS from Pacific Biosciences (PacBio), Bionano Genomics, and Strand-seq technology (Chaisson et al., 2019; Ebert et al., 2021); however, this is the first time that nearly all 1kGP trios have been sequenced at high coverage and jointly analyzed for the discovery and genotyping of genomic variation across the size and frequency spectrum, ranging from SNVs to large and complex SVs in a singular resource. We sequenced the expanded cohort of 3,202 samples to a targeted depth of 30X (minimum 27X, mean 34X) genome coverage using Illumina NovaSeq 6000 instruments. We aligned reads to the GRCh38 reference and performed SNV/INDEL calling using GATK’s HaplotypeCaller. Using this strategy, we called over 111 million SNVs and over 14 million INDELs with false discovery rate (FDR) of 0.3% and 1.15%, respectively, across the entire cohort of 3,202 samples. We also discovered and genotyped a comprehensive set of SVs, including insertions, deletions, duplications, inversions, and multiallelic copy number variants, by integrating multiple algorithms and analytic pipelines, including GATK-SV (Collins et al., 2020), svtools (Larson et al., 2019), and Absinthe (Corvelo et al., 2021). Comparison with previous low coverage sequencing performed in phase 3 of the 1kGP demonstrates significant improvements in sensitivity and specificity in the SNV, INDEL and SV call sets, highlighting that the re-sequencing effort and expansion of the cohort to include trios brought significant value to the resource.

One of the major applications of the phase 3 1kGP call set has been its widespread use as a reference panel for variant imputation in sparse, array-based genotyping data with a goal of improving the statistical power of downstream genome-wide association studies (GWAS) and facilitating fine-mapping of causal variants. We leveraged the presence of full trios in the expanded 1kGP cohort and performed haplotype phasing of SNVs and INDELs using a statistical phasing approach with pedigree-based correction. We demonstrate the improvements brought by including family members when phasing variants and show how it compares to the phase 3 version. Finally, we evaluate the imputation performance of the high coverage panel and demonstrate improved results especially in INDEL imputation as compared to the phase 3 panel.

Over the past few years, the cost of high coverage WGS has decreased dramatically which, combined with substantial progress in analytics tools, has contributed to the emergence of several population-scale high coverage WGS panels, such as the Genome Aggregation Database (gnomAD; 76,156 WGS and 125,748 WES samples) (Karczewski et al., 2020), Trans-Omics for Precision Medicine (TOPMed, ∼180,000 samples) (Taliun et al., 2021), or the UK Biobank (UKBB, goal to sequence 500,000 samples by 2023) to name a few. These growing resources, many fold larger in sample size than the 1kGP cohort, enable continuous expansion of the catalog of genetic variation in the human population and facilitate discoveries that improve human health. Unlike the 1kGP, these recent large-scale WGS efforts have restrictions on public data sharing as they are often linked to clinical data, which amplifies privacy concerns. As a result, only aggregate population-level allele frequencies are typically available for public access. In contrast, samples within the 1kGP cohort have been consented for full public release of genetic information which allows for unrestricted sharing of the complete sample-level genotype (GT) data. This enables granting access to a downloadable reference imputation panel, as well as use of the dataset for methods development and benchmarking, among other applications. All the data generated as part of this project, including CRAM files and VCFs, have been made publicly available (see the Resource Availability section). We envision this updated version of the 1kGP cohort to become the new de facto public resource for the worldwide scientific community working on genomics and genetics.

## RESULTS

### Small variation across the 3,202 1kGP samples

Using the Illumina NovaSeq 6000 System, we performed WGS of the original 2,504 1kGP unrelated samples as well as additional 698 related samples. This completed 602 parent-child trios in the 1kGP cohort and brought the total number of sequenced and jointly-genotyped samples to 3,202 (Figure 1A, Table S1). At the cohort-level, we discovered a total of 117,256,690 small variant loci, which represent 125,484,020 unique alternate alleles, including 111,048,944 SNVs and 14,435,076 INDELs (Table 1). Across all SNVs and INDELs, there are 58,379,163 (47.6%) singletons (allele count (AC)=1), 45,931,977 (37.5%) rare (allele frequency (AF) ≤ 1%), and 18,212,589 (14.9%) common (AF > 1%) alleles, as defined using AF estimates based on unrelated samples in the cohort (Figure 1B). Out of all small variants, 19,237,848 (15.3%) represent novel alleles, defined here as not reported in dbSNP build 151 (Sherry et al., 1999). Among the novel variants, 63.0% are singletons, 32.3% are rare, and 4.7% are common in the population (Figure 1B).

**Figure 1.**
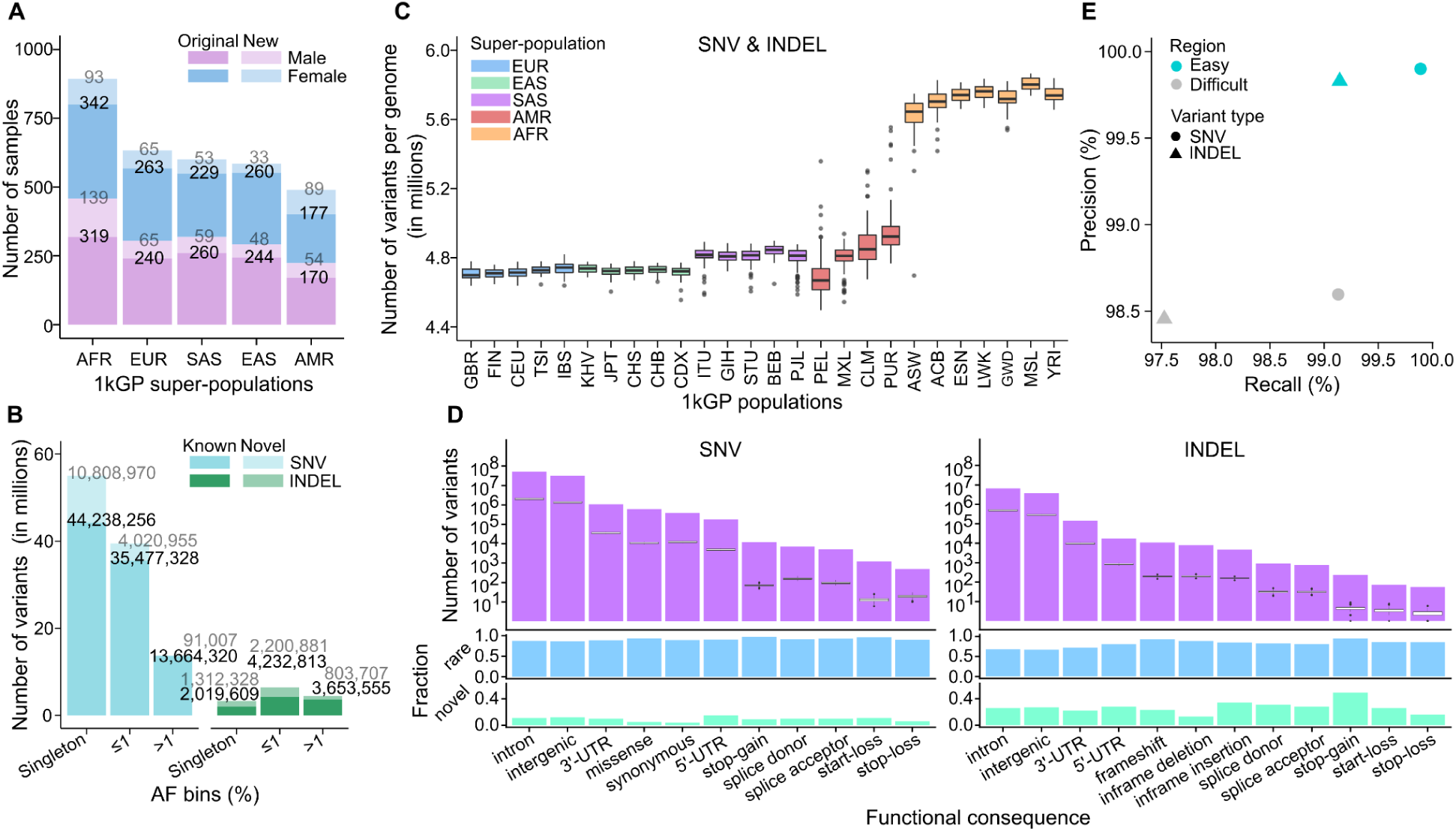
SNV/INDEL discovery in the high coverage WGS data across the 3,202 1kGP samples. **(A)** Counts of sequenced 1kGP samples stratified by sex and super-population. Transparent areas represent counts coming from the newly added 698 related samples that were not part of the original 1kGP call set. **(B)** Cohort-level counts across the 3,202 samples, stratified by variant type, AF bins, and novel/known variants (absent from/present in dbSNP build 151). AF was estimated based on the 2,504 unrelated samples only. Singleton counts are reported as a separate bin and were excluded from the ≤1% AF bin. **(C)** Total number of small variant loci per genome, stratified by population. Counts are restricted to variants that passed VQSR. See also Figure S1A-C. **(D)** Predicted functional SNVs and INDELs. Top row: cohort-level counts (purple bar plot) overlaid with distributions of per-genome counts (box plots) across the 2,504 unrelated samples. Middle row: fraction of rare (MAF ≤ 1%) SNVs and INDELs among the predicted functional sites. Bottom row: fraction of novel (defined above) SNVs and INDELs among the predicted functional sites. See also Figure S1G, S1H. **(E)** Precision vs. recall of SNVs and INDELs computed relative to the GIAB NA12878 truth set v3.3.2, stratified by easy and difficult regions of the genome. See also Figure S1D. Super-population ancestry labels: European (EUR), African (AFR), East Asian (EAS), South Asian (SAS), American (AMR). For descriptions of population labels please refer to Table S1.

**Table 1.** Summary of variant counts in the high coverage 1kGP call set at the cohort and sample level. Super-population ancestry labels: European (EUR), African (AFR), East Asian (EAS), South Asian (SAS), American (AMR). SV types: DEL: deletion, DUP: duplication, mCNV: multiallelic copy number variant, INS: insertion, INV: inversion, CPX: complex SV, CTX: inter-chromosomal translocation.

| Variant type | # variants across 3,202 samples | Average # variants per sample |  |  |  |  |  |
| --- | --- | --- | --- | --- | --- | --- | --- |
|  | Total | All | AFR | EUR | SAS | EAS | AMR |
| **Total small variants** | **125,484,020** | **4,952,915** | **5,623,313** | **4,645,189** | **4,736,023** | **4,651,279** | **4,754,817** |
| SNV | 111,048,944 | 4,080,992 | 4,653,521 | 3,818,951 | 3,896,324 | 3,822,328 | 3,911,413 |
| INDEL-DEL | 8,010,854 | 451,277 | 503,995 | 426,940 | 433,635 | 428,078 | 435,976 |
| INDEL-INS | 6,424,222 | 420,646 | 465,797 | 399,298 | 406,064 | 400,873 | 407,428 |
| **Total SVs** | **173,366** | **9,679** | **10,742** | **9,176** | **9,304** | **9,251** | **9,363** |
| DEL | 90,259 | 4,232 | 4,715 | 4,001 | 4,066 | 4,035 | 4,089 |
| DUP | 28,242 | 1,207 | 1,322 | 1,153 | 1,168 | 1,155 | 1,178 |
| mCNV | 673 | 425 | 433 | 422 | 419 | 425 | 419 |
| INS | 49,693 | 3,534 | 3,971 | 3,329 | 3,378 | 3,361 | 3,403 |
| INV | 920 | 68 | 71 | 66 | 67 | 67 | 66 |
| CPX | 3,568 | 213 | 230 | 205 | 206 | 208 | 208 |
| CTX | 11 | 0 | 0 | 0 | 0 | 0 | 0 |

To better characterize our variant calls, we divided the genome into easy- and difficult-to-sequence regions (see Methods), based on stratification intervals generated by the Genome in a Bottle (GIAB) Consortium (Krusche et al., 2019). At the locus-level, 6.6% (7,706,051 out of 117,256,690) of small variant sites across the 3,202-sample cohort are multiallelic. The difficult regions constitute only 20% of the genome but they contain a disproportionately high fraction of multiallelic sites (74.5%) as well as INDEL loci (64.3%). This is in contrast to SNV loci which fall mostly in the easy regions of the genome (easy: 76.9% vs. difficult: 23.1%). The enrichment for multiallelic and INDEL calls in difficult regions is consistent with expectation, as these regions mostly consist of low complexity and repetitive elements where alignment of sequencing reads is particularly challenging and where INDELs are known to typically form (Montgomery et al., 2013).

At a genome level, we called an average of 4,952,915 small variants (Figure 1C, Table 1). This includes an average of 4,080,992 SNVs and 871,923 INDELs per genome, across samples from all populations (Table 1, Figure S1A, S1B). We observed an average transition to transversion ratio (Ti/Tv) of 2.01 and heterozygous to non-reference homozygous ratio (Het/Hom) of 1.70 (Figure S1C), consistent with expectations for WGS data. As expected, the average number of variant sites was higher in the individuals from populations with African ancestry (AFR), with 4,653,521 SNVs and 969,792 INDELs per genome (Table 1). In line with that, we also observed a higher Het/Hom ratio of 2.03 among the AFR samples (Figure S1C). We also noticed higher variability in the number of variants in individuals belonging to the admixed populations with American ancestry (AMR) (Figure 1C).

### Predicted functional consequence of small variants

To assess functional consequences of SNVs and INDELs in the high coverage call set, we annotated all variant loci using the Ensembl Variant Effect Predictor (VEP) v104 tool (McLaren et al., 2016). Across the unrelated samples, we found a total of 605,896 missense, 384,451 synonymous, as well as 36,520 predicted loss of function mutations (pLOF), defined here as stop-gains (n=12,181), frameshift (n=10,850), and splice mutations (n=13,489) (Figure 1D, purple bar plots). Depending on the functional consequence category, 86-97% and 67-95% of predicted functional SNVs and INDELs, respectively, are rare (MAF ≤ 1%), with 4-15% SNVs and 13-49% INDELs being novel (*i.e.* absent from dbSNP build 151) (Figure 1D, blue and green bar plots). At a genome level, we found on average 10,625 missense, 11,956 synonymous, 76 stop-gain, 195 frameshift, and 314 splice mutations, without applying MAF filtering (Figure 1D, Table S3). At MAF ≤ 1%, each sample carries on average 11 stop-gain, 18 essential splice, and 14 frameshift mutations. These cohort- and genome-level counts are consistent with previous reports (Karczewski et al., 2020; Taliun et al., 2021). As expected, AFR samples carry the highest counts of variants across all functional categories as compared to other populations (Figure S1G, Table S3), with magnitudes of difference between populations being similar across high and low impact functional categories (Figure S1H).

### False discovery rate among small variants

We determined the FDR of the high coverage call set by comparing our genotype calls in sample NA12878 to the GIAB NA12878 truth set v3.3.2 (Zook et al., 2019), in the high confidence regions of the genome. Using this approach, the estimated FDR is 0.3% for SNVs and 1.15% for INDELs. We observed ∼10-fold lower FDR (=1-Precision in Figure 1E) in the easy as compared to difficult subsets of the high confidence regions for both SNVs (0.10% vs. 1.40%, respectively) and INDELs (0.17% vs. 1.54%) (Figure 1E & S1D). In the easy regions, sensitivity of SNV and INDEL calls reaches 99.89% and 99.14%, respectively, whereas in difficult regions it is slightly lower, 99.13% for SNVs and 97.53% for INDELs (Figure 1E & S1D).

To further evaluate the quality of the high coverage call set, we separately analyzed the subset of small variants which tends to be the most enriched for false positive calls, namely the singletons (defined here as variants with AC=1 across the entire 3,202-sample cohort). Due to the mixed nature of the expanded 1kGP cohort, which now includes both unrelated as well as related samples, it is important to note that the number of singletons per genome varies depending on the sample’s relatedness status, with children having the fewest singletons, followed by parents, and unrelated samples in the cohort (Figure S1E). We observed a nearly bimodal distribution of per-genome singleton counts among children with modes at 444 and 1,108 and the mean of 1,340, and a unimodal distribution among parents as well as unrelated samples with means at 12,365 and 23,197, respectively (Figure S1E). These differences are due to “private” variants (*i.e.* inherited variants that are private to a single family) which are not being counted as singletons in children, while 50% and 100% of them are being counted as singletons in each of the parents and in unrelated samples, respectively. Singletons among children that are part of the 602 trios in the cohort represent putative *de novo* mutations (DNMs). The expected number of germline DNMs is ∼100 per child (Jónsson et al., 2017; Kong et al., 2012), which suggests that the mean number of singletons among children exceeds the expectation by about a factor of 10, although this varies rather widely from sample to sample (Figure S1E). Given that all 1kGP samples are derived from lymphoblastoid cell lines (LCLs) of various ages, these additional singletons likely represent somatic artifacts from cell line propagation (Ng et al., 2021), as well as some false positive calls. As evidence of the presence of somatic artifacts, we observed aneuploidy of allosomes in 0.94% of the samples, and sub-chromosomal level mosaic copy number variants (CNVs) on multiple autosomes (Figure S2). This is in agreement with findings from the Polaris project (Illumina Inc., 2019).

To estimate FDR among singletons, we evaluated singletons in sample NA12878 against the GIAB NA12878 truth set v3.3.2 (Zook et al., 2019). Since NA12878 is a child in the expanded 1kGP cohort, we jointly assessed both its *de novo* variants (n=2,404) as well as inherited heterozygous variants that are private to the NA12878 trio (n=15,131). Based on that, the estimated FDR among singletons is 1.01% (see Methods; Figure S1F). To ensure that this approach for FDR estimation is not biased due to inclusion of NA12878’s parents in joint-genotyping, we also computed FDR among singletons in NA12878 from an independent jointly-genotyped high coverage call set consisting of just the original 2,504 1kGP unrelated samples. Using this orthogonal validation, the estimated FDR is 0.98%, which agrees closely with the analysis based on the 3,202-sample call set (Figure S1F). Additionally, we evaluated singletons against the recently released GIAB NA12878 truth set v4.2.1 (Wagner et al., 2021). Thanks to inclusion of additional technologies such as PacBio-HiFi and 10X Genomics, the GIAB v4.2.1 truth set excludes some of the calls believed to be mosaic variants that arose due to cell line propagation that were present in the GIAB v3.3.2 truth set. Based on this comparison, the FDR among singletons is 5.93% (combined analysis of DNMs and private variants in NA12878 from the 3,202-sample joint call set) or 5.78% (analysis of singletons from the 2,504-sample joint call set) (Figure S1F; see Methods). This indicates that ∼5% of singleton calls in the high coverage call set appear to be truly present in the cell lines but may not represent true population variants or even real DNMs in the original donors, highlighting potential shortcomings of using cell line derived DNA for this study.

### Structural variation across the 3,202 1kGP samples

We generated an SV call set across all 3,202 1kGP samples with short read sequencing data. These SV genotypes were discovered and integrated from three analytic pipelines: GATK-SV (Collins et al., 2020), svtools (Abel et al., 2020), and Absinthe (Corvelo et al., 2021) (see Methods) (Table S4, Figure S3). This final ensemble call set included 173,366 loci, comprised of 90,259 deletions, 28,242 duplications, 673 multiallelic copy number variants (mCNVs), 49,693 insertions, 920 inversions, 3,568 complex SVs (CPX) consisting of a combination of multiple SV signatures, and 11 inter-chromosomal translocations (CTX, Figure 2A, Table 1). The size and allele frequency distribution of SVs followed expectations; mobile element signatures were observed for ALU (200-300 bp), SVA (1-2 kb), and LINE1 (5-6 kb) variants (Figure 2B). Most SVs were rare, and SV allele frequencies were inversely correlated with SV size (Figure 2C). On average, ∼9,679 SVs were discovered in each genome (see Figure 2D, Table 1). The distribution of SVs observed per individual followed expectations for ancestry with the greatest number of SVs per genome derived from AFR populations (Figure 2E, Table 1) (Campbell and Tishkoff, 2008). The specificity of the SV call set was also quite high, with a *de novo* SV rate of 3.2%, which represents the combination of false positive SVs in children, false negative SVs in parents, and cell line artifacts in children (Figure 2F).

**Figure 2.**
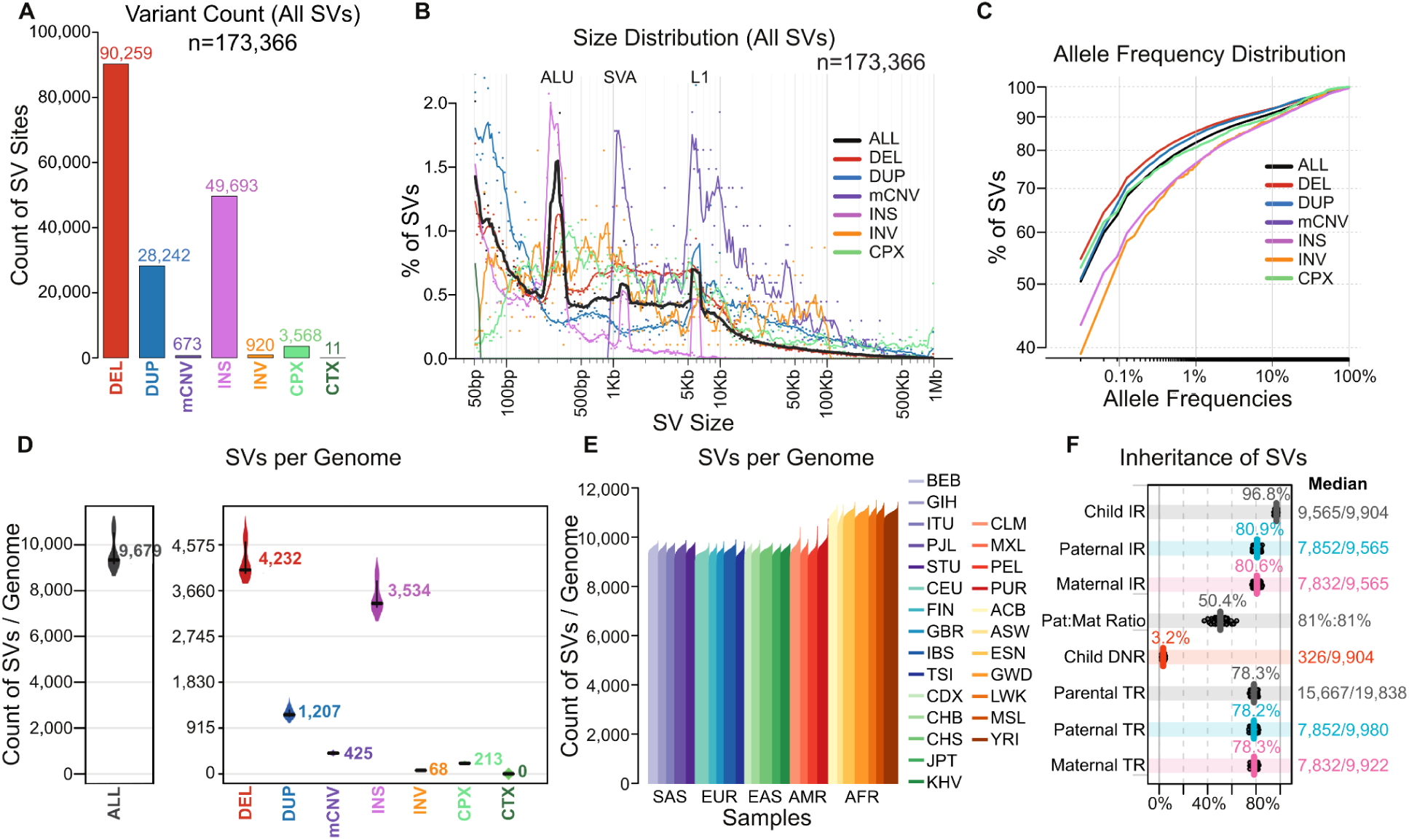
SV discovery in the high coverage WGS data across the 3,202 1kGP samples. **(A)** The count, **(B)** size distribution, and **(C)** allele frequency distribution of each SV for each SV class is shown. The mean per sample count of SVs by variant class **(D)** and ancestral population **(E)** is also provided, as well as **(F)** inheritance rates of all SVs. In (F) paternal and maternal inheritance rates (IR) refer to the proportions of SVs in children’s genome that are paternally and maternally inherited, and parental transmission rate (TR) refers to the proportion of SVs in parents’ genomes that are transmitted. DNR: *De Novo* Rate. SV types: DEL: deletion, DUP: duplication, mCNV: multiallelic copy number variant, INS: insertion, INV: inversion, CPX: complex SV, CTX: inter-chromosomal translocation. Super-population ancestry labels: European (EUR), African (AFR), East Asian (EAS), South Asian (SAS), American (AMR). For descriptions of population labels please refer to Table S1. See also Figure S3.

### Comparison of the small variant calls to the 1kGP phase 3 call set

To quantify the improvements that the high coverage sequencing and pipeline upgrades brought to the new 1kGP call set, we compared our small variant calls against the original phase 3 call set. For consistency, we restricted this comparison to variants discovered in the 2,504 samples that are common to both datasets. For that purpose, we generated an independent jointly-genotyped high coverage call set, consisting of just the original samples, and used it for the comparison. Direct comparison to the original call set was not possible as the phase 3 dataset was aligned to the GRCh37 reference. To overcome this issue, we lifted-over the phase 3 call set to the GRCh38 build using CrossMap (Zhao et al., 2014) (see Methods).

The 2,504-sample high coverage call set includes 96,950,998 SNVs and 13,132,415 INDELs across the autosomes. This represents a 1.24-fold cohort-level increase in the number of SNVs and 4.05-fold increase in the number of INDELs, compared to the lifted-over phase 3 call set (78,324,761 SNVs and 3,244,241 INDELs). Among SNVs, we observed the largest gains in the number of singletons (AC=1; gain of 15,123,906 SNVs) and rare (AC > 1 and AF ≤ 1%; gain of 3,557,925 SNVs) allele categories in the high coverage relative to the phase 3 call set. As expected, the number of common (AF > 1%) SNVs was similar across both call sets (Figure 3A). In the case of INDELs, we observed gains across the entire AF spectrum relative to the phase 3 call set (Figure 3B). The highest increase (676-fold) is in the singleton category where the phase 3 call set contains only 4,437 singleton INDELs, as compared to 2,999,027 in the high coverage call set. The low number of ultra-rare INDEL calls in the phase 3 set can be attributed to more stringent filtering applied to INDELs as compared to biallelic SNVs (The 1000 Genomes Project Consortium, 2015) and limitations of low coverage sequencing. The increase in the number of rare and common INDELs in the high coverage vs. phase 3 call set is also significant, 3.49- and 2.72-fold, respectively (Figure 3B). Additionally, we called significantly more INDELs above 50 bp in length (182,579 vs. 2,172 in phase 3, Figure S4A).

**Figure 3.**
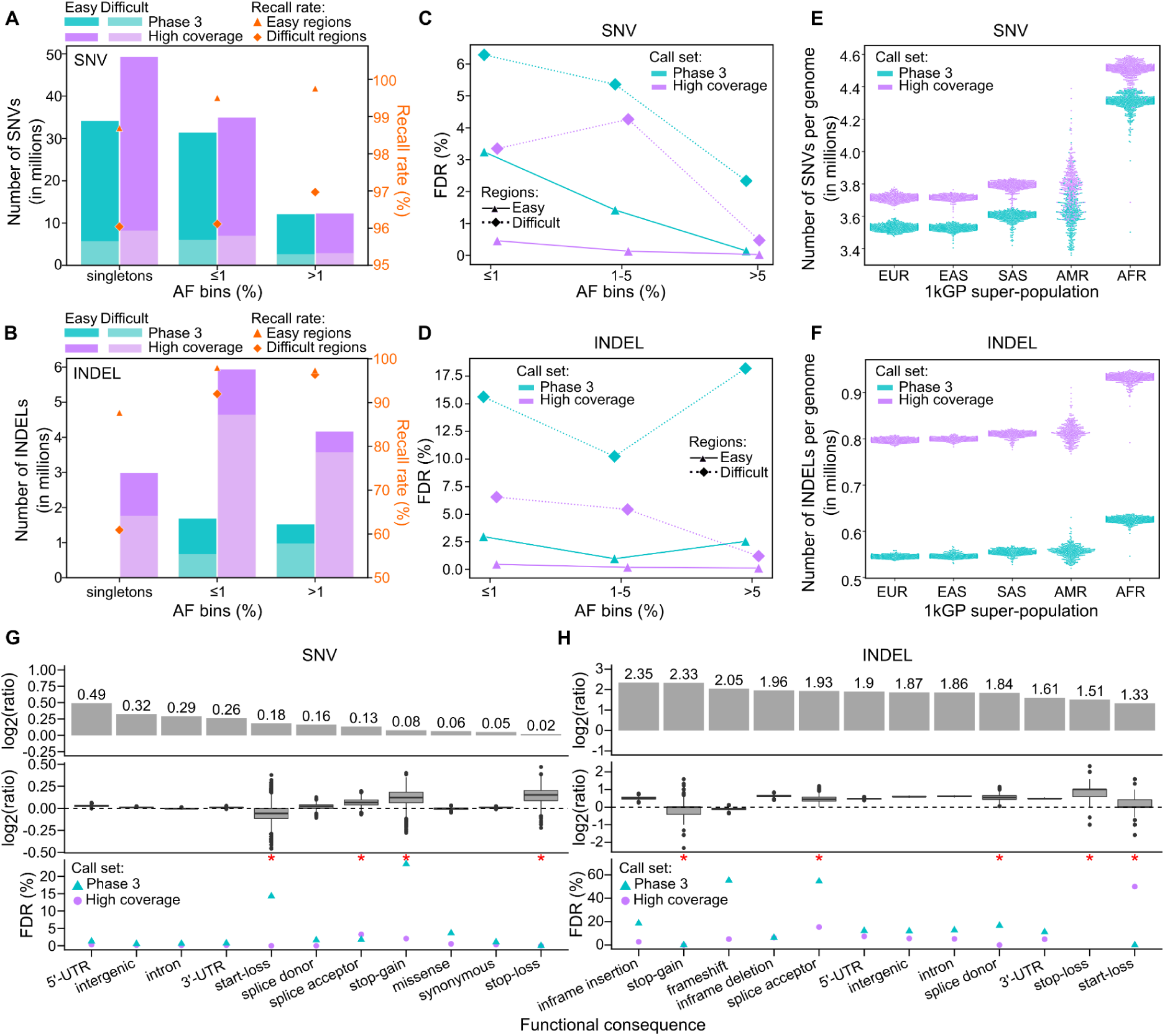
Comparison of small variant calls to the phase 3 call set. Number of autosomal SNVs **(A)** and INDELs **(B)** across the 2,504 samples in phase 3 and high coverage datasets, stratified by AF bins. Counts of variants with AC=1 (singletons) are reported as a separate bin and were excluded from the ≤1% AF bin. Multiallelic loci were split into separate lines and INDEL representation was normalized prior to counting. Secondary y axis: % of autosomal phase 3 variants recalled in the high coverage call set across SNVs (A) and INDELs (B) in easy and difficult regions of the genome. See also Figure S4C. Comparison of FDR across SNVs **(C)** and INDELs **(D)** between the high coverage and phase 3 call sets, stratified by regions of the genome. See also Figure S4B. Genome-level SNV **(E)** and INDEL **(F)** counts in the phase 3 vs. high coverage call sets, stratified by 1kGP super-population ancestry (EUR: European, AFR: African, EAS: East Asian, SAS: South Asian, AMR: American). Reported counts are at a locus level. Comparison of predicted functional SNV **(G)** and INDEL **(H)** counts in the high coverage vs. phase 3 call set. Log2(ratio) denotes log2(ratio of variant counts in the high coverage vs. phase 3 call set). Top row: cohort-level comparison. Middle row: genome-level comparison. Bottom row: comparison of FDR. Red asterisks on top of the FDR plot denote categories with fewer than 100 sites in sample NA12878 (*i.e.* categories where FDR estimation is less reliable). See also Figure S4D,E. FDR in panels C, D, G, and H was estimated based on comparison of calls in sample NA12878 to the GIAB NA12878 truth set v3.3.2.

Overall, we recalled 98.3% of the phase 3 small variants in the high coverage call set. SNV recall rate is over 99% in the easy regions of the genome, and over 96% in the difficult regions across all AF bins (Figure 3A). In the easy regions of the genome, INDEL recall rate is above 97% in rare and common AF bins, but is down to 88% among singletons (Figure 3B). In the difficult regions of the genome, INDEL recall rate decreases significantly with decreasing AF, from 96% in the common allele bin, 92% in the rare, down to only 61% in the singleton bin (Figure 3B). We observed high correlation of AF among shared variants between the high coverage and phase 3 call sets (Figure S4C). In the easy regions of the genome, the Pearson correlation coefficient of AFs is above 0.99 for both SNVs and INDELs, whereas in the difficult regions it drops to 0.98 for SNVs and 0.93 for INDELs (Figure S4C).

At a per sample level, we observed a 1.05-fold average increase in the number of SNVs and 1.47-fold increase in the number of INDELs in the high coverage (3,950,455 SNVs and 838,249 INDELs across the 2,504 samples) as compared to the phase 3 call set (3,759,570 SNVs and 570,067 INDELs; Figure 3E, 3F).

The FDR of the 2,504-sample high coverage call set is 0.1% for SNVs and 1.1% for INDELs, as compared to 0.6% for SNVs and 12.4% for INDELs in the lifted-over phase 3 call set. When we stratified the FDR estimation by AF bins and genomic regions, we observed significantly lower FDR across the entire AF spectrum, in both easy and difficult genomic regions, in the high coverage as compared to the phase 3 call set (Figure 3C, 3D, S4B). This trend was particularly pronounced among rare (AF ≤ 1%) SNVs and INDELs in the difficult regions of the genome.

We observed 1.01-1.40-fold increase in the number of SNVs falling into various functional categories in the high coverage as compared to the phase 3 call set at a cohort-level (Figure 3G, top row). This increase was especially pronounced (≥ 1.2-fold) in the intronic, intergenic, and untranslated region (UTR) categories. The rather insignificant increase in the number of SNVs in protein-coding categories (1.01-1.13-fold; Figure 3G, top row) was expected since variant discovery in the phase 3 call set was based on high coverage WES, in addition to low coverage WGS. We observed a more significant increase, between 2.5- and 5-fold, in the number of predicted functional INDELs in the high coverage vs. phase 3 call set at the cohort-level (Figure 3H, top row). This is consistent with larger overall gains in INDEL calls as compared to SNVs in the high coverage call set. At a genome level, the ratios of predicted functional SNV counts in the high coverage vs. phase 3 call set were close to 1 (*i.e.* no significant difference) with well-controlled FDR in both call sets across nearly all categories. The two exceptions were stop-loss and stop-gain categories for which mean ratios were 1.11 and 1.09 (Figure 3G, middle row), respectively, with stop-gain category having a particularly high FDR in the phase 3 call set (25% vs. 2% in high coverage; Figure 3G, bottom row; Figure S4D). Among INDELs, the genome-level ratios in high coverage vs. phase 3 were higher than for SNVs, reaching over 1.5 in case of inframe deletions, as well as intergenic, and intronic INDELs, consistently with larger proportion of common INDELs relative to SNVs among new loci discovered in the high coverage call set. We observed a slight decrease in the mean number of frameshift (7%) and stop-gain (11%) INDELs in the high coverage as compared to the phase 3 call set (Figure 3H, middle row). In case of frameshift mutations, this decrease might be explained by a particularly high FDR rate in the phase 3 call set (55% vs. 5% in the high coverage call set; Figure 3H bottom row; Figure S4E).

### Comparison of the SV calls to the 1kGP phase 3 call set

The ensemble SV call set was compared to the 1kGP phase 3 SVs (7.4X average coverage) (Sudmant et al., 2015) on the 2,504 shared samples to assess the quality and unique value brought by high coverage sequencing and genotyping capabilities of new analytic pipelines. The current ensemble SV call set discovered over two-fold more SV sites than phase 3 (169,713 vs. 68,697), and encompassed 87.7% of the phase 3 SV calls (Figure 4A). This increased sensitivity and high overlap of phase 3 SVs was consistent across all SV classes (Figure 4A), with an average of 9,655 SVs detected per genome in the current ensemble call set compared to 3,431 SVs in the phase 3 call set (Figure 4B).

**Figure 4.**
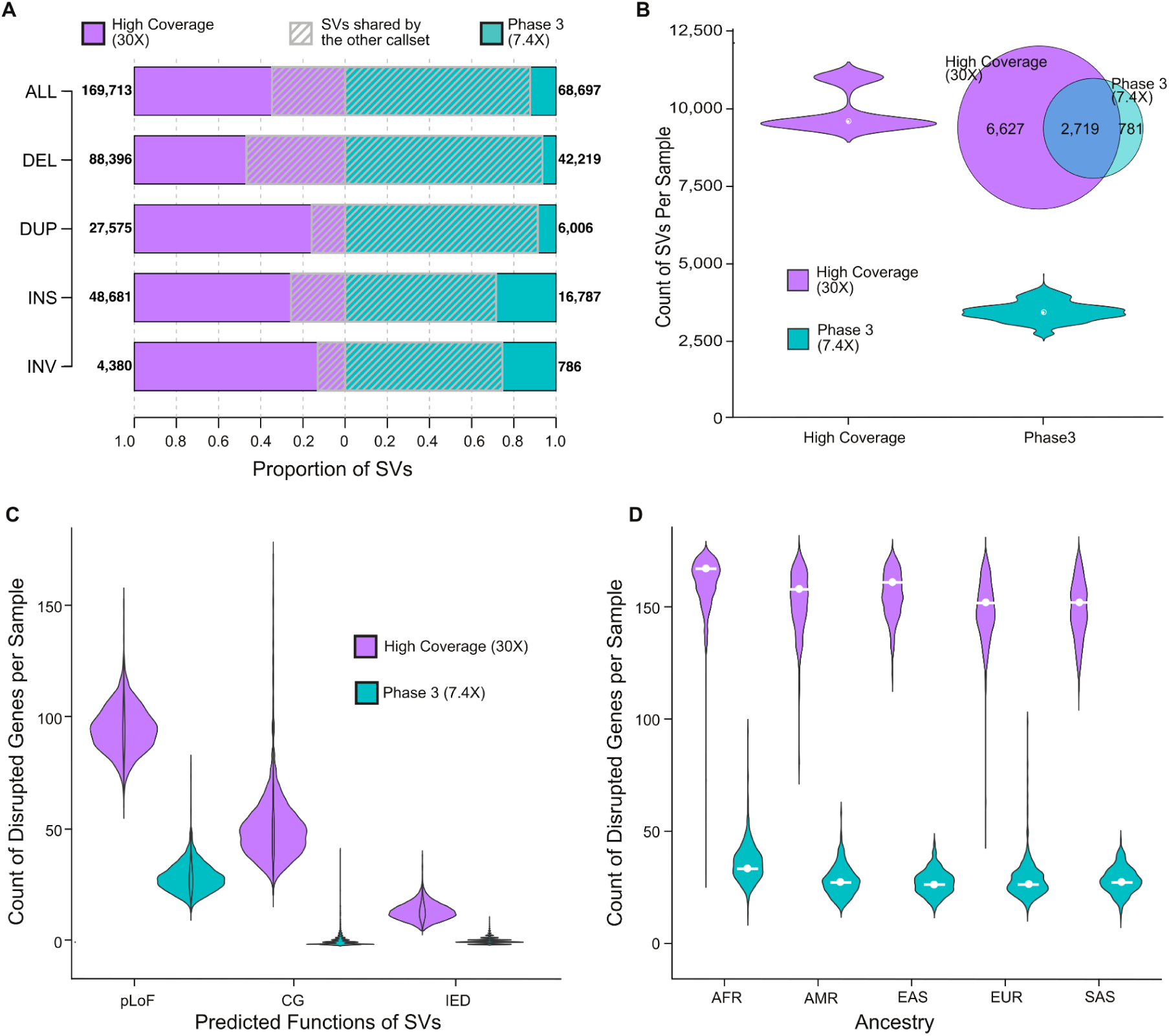
Comparison of the ensemble SV calls to the phase 3 call set. **(A)** Count of SV sites in the current ensemble SV call set and low coverage phase 3 SV call set, and their overlap. Numbers next to each bar represent the counts of SV sites in each dataset. **(B)** The distribution of SV counts per sample in both call sets and their average overlap, displayed in the Venn diagram. **(C)** Count of genes altered by SVs in both datasets. pLoF: probable loss of function, CG: complete copy gain, IED: intragenic exon duplication. **(D)** Count of genes altered by SVs across ancestral populations. See also Figure S5.

The high coverage SV call set provided significant added value in terms of the discovery of SVs that alter gene function by comparison to the phase 3 low-coverage SV dataset. Consistent with a previous large population study from short-read WGS that predicted disruption of 180 genes by SVs in each genome (Collins et al., 2020), as well as a recent study from the HGSVC using long-read WGS and complementary technologies that estimated 189 SVs per genome that altered protein coding genes (Ebert et al., 2021), we observed that biallelic SVs in each genome resulted in alteration of 162 genes per genome, including probable loss of function (pLoF) of 97 protein coding genes, complete copy gain (CG) of 50 genes, and duplications of intragenic exons (IED) of 15 genes. Notably, the functional impact of IEDs has been previously shown to be correlated with negative selection against LoF point mutations (Collins et al., 2020). This represents a considerable increase in the estimates from the low-coverage phase 3 call set that predicted an average of 32 genes disrupted by SVs per genome (30 pLoF, 1 CG and 1 IEDs; Figure 4C, S5). The high-coverage 1kGP dataset also offered an estimate in the population diversity of functional SV variation, where AFR populations had the highest number of SVs per genome (Figure 4D), and similar patterns were observed when evaluating pLoF, CG and IED SVs that altered protein coding gene sequences individually (Figure S5).

### Haplotype phasing of small variants

We performed haplotype phasing of high quality SNVs and INDELs across the 3,202-sample 1kGP cohort using statistical phasing with pedigree-based correction (chromosome X was phased without the correction; see Methods). Prior to phasing, we filtered the small variant call set (see Methods) which resulted in a selection of 72,065,314 high quality variants across autosomes and chromosome X (Figure 5A). Included among these are 61,411,215 SNVs, 9,954,481 INDELs, and 699,618 multi-nucleotide polymorphisms (MNPs) (counts at the alternate allele (ALT) level) (Table S6).

**Figure 5.**
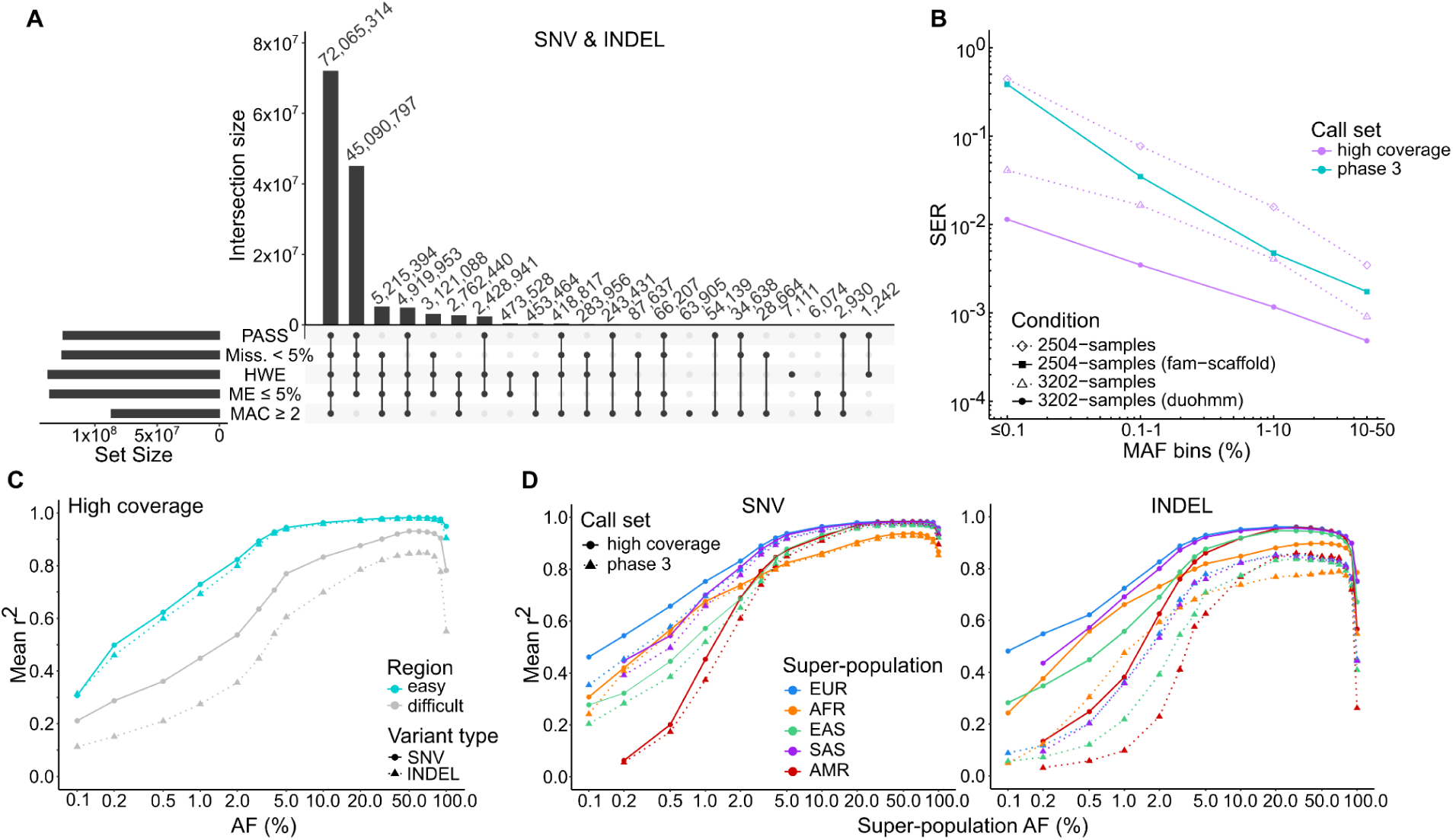
Small variant phasing and imputation performance. **(A)** Counts of small variants passing specified filtering criteria. Multiallelic variants were split into separate rows prior to counting. PASS: sites that passed VQSR; Miss.: genotype-level missingness; HWE: HWE exact test p-value > 1e-10 in at least one of the five 1kGP super-populations; ME: mendelian error rate across complete trios; MAC: minor allele count. See also Table S6. **(B)** Haplotype phasing accuracy of the high coverage (hc) and the phase 3 (ph3) 1kGP panel. SER: switch error rate in sample NA12878 relative to the Platinum Genome truth set (autosomes only). Conditions: (1) 3,202-sample (duohmm): hc panel phased using statistical phasing with pedigree-based correction; (2) 3,202-samples: hc panel phased using statistical phasing without the pedigree-based correction; (3) 2,504-samples: hc panel phased using statistical phasing alone (unrelated samples only). (4) 2,504-samples (fam-scaffold): ph3 panel phased using statistical phasing with family-based scaffold (unrelated samples only). The two hc panels represented with dashed lines were created for evaluation purposes only. See also Figure S6A,B. **(C)** Imputation accuracy of SNV and INDEL genotypes imputed using the high coverage panel, stratified by genomic regions. Mean r^2^: squared Pearson correlation coefficient between imputed allelic dosages and dosages from WGS data, averaged over 110 Simons Genome Diversity Project samples. See also Figure S6C, S6D. **(D)** Comparison of the imputation accuracy between the high coverage and phase 3 panels for SNVs and INDELs, stratified by super-population ancestry (EUR: European, AFR: African, EAS: East Asian, SAS: South Asian, AMR: American). This comparison was restricted to sites that are shared between the two panels.

We evaluated phasing accuracy of the phased panel by computing switch error rate (SER) in sample NA12878 relative to the gold standard phasing truth set, *i.e.* Platinum Genome NA12878 call set generated by Illumina (Eberle et al., 2017). The SER across all autosomes was 0.074% (1,754 switch errors (SEs) across 2,338,955 assessed SNV/INDEL heterozygous (HET) pairs), indicating high accuracy of phasing. As expected, chromosome X (phased using statistical phasing alone) displayed higher SER as compared to autosomes (SER=0.491%; 362 SEs across 73,794 HET pairs; Figure S6A). We did not observe a significant difference in phasing accuracy between SNVs and INDELs, other than on chromosome X, where SER for INDELs was 2.01% (187 SEs across 9,298 assessed INDEL HET pairs) as compared to 0.51% for SNVs (328 SEs across 64,583 SNV HET pairs) (Figure S6B). We observed an expected increase in SER with decrease in MAF, but the SER remained low throughout the entire MAF spectrum, reaching a maximum of 1.14% in the ≤ 0.1% MAF bin across autosomes (Figure 5B, violet solid line; see Figure S6A for per chromosome breakdown). Such high phasing accuracy at the low end of the MAF spectrum can be attributed to both the presence of family members in the expanded 1kGP cohort (Figure 5B, dashed violet line with open triangles vs. dashed violet line with open diamonds) as well as pedigree-based correction applied after statistical phasing (Figure 5B, solid violet line vs. dashed violet line with open triangles).

Finally, we compared the phasing accuracy of the high coverage panel to the phase 3 panel, which was phased using statistical phasing with family-based scaffold built from chip array data (The 1000 Genomes Project Consortium, 2015). The overall SER across autosomal SNVs and INDELs for the phase 3 panel was 0.76% (16,938 SEs across 2,238,400 HET pairs), which is 10-fold higher than SER in the high coverage call set. The SER on chrX was 1.29% (879 SEs across 68,290 HET pairs) in the phase 3 panel, which is 2.6-fold higher than SER on chrX in the high coverage panel. The significantly lower SER in the high coverage as compared to the phase 3 panel was also observed in a stratified analysis across all four MAF bins (Figure 5B, solid violet vs. solid aqua line), with magnitude of decrease ranging from 3.6-fold in the case of the most common MAF bin up to 34-fold in the rarest MAF bin. The significant improvement in phasing accuracy underscores the benefit of including trios while building the new panel. It is worth noting that the phasing accuracy of the 2,504-sample phase 3 dataset was slightly better than that of the unrelated 2,504-sample high coverage dataset (Figure 5B, solid aqua line vs. dashed violet line with open diamonds) due to the fact that the latter dataset was phased using statistical phasing alone, without the family-based scaffold.

### Imputation performance of the small variant reference panel

To assess imputation performance of the high coverage panel, and to compare it with the original phase 3 panel, we imputed a set of 279 diverse samples from the Simons Genome Diversity Project (SGDP) (Mallick et al., 2016) with either the high coverage or the phase 3 panel as the reference. We evaluated the accuracy of imputed genotypes by computing the squared Pearson correlation coefficient (r^2^) between imputed allelic dosages and dosages from publicly available WGS data (Mallick et al., 2016) used here as a truth set (see Methods). Using the high coverage reference panel, we observed significantly higher mean imputation accuracy, for both SNVs and INDELs, in easy as compared to the difficult regions of the genome (r^2^=0.8 attained at AF=∼2% for SNVs and INDELs in easy regions, compared to AF=∼10% and AF=∼30% for SNVs and INDELs, respectively, in difficult regions) (Figure 5C). This expected pattern of improved imputation performance in easy-to-sequence regions was also observed in a stratified analysis, across all five super-populations (Figures S6C, S6D).

When compared with the phase 3 reference panel, the high coverage panel displays superior imputation accuracy across shared loci for all five super-populations, especially in the case of INDELs and, to a lesser extent, rare SNVs (Figure 5D). Depending on the ancestry group, the high coverage panel achieves a mean imputation accuracy r^2^ of 0.8 at AF=1-4% across SNVs and AF=2-4% across INDELs (Figure 5D). In the case of the phase 3 panel, the mean r^2^=0.8 is achieved at AF=2-4% across SNVs and AF=5-70% (with AFR super-population being the biggest outlier at AF=70%) across INDELs (Figure 5D). Improvements in INDEL imputation using the high coverage panel are significant across the entire AF spectrum. At an AF=0.5%, for example, we observe a mean r^2^ increase ranging from 1.8-fold (AFR) to 4.3-fold (AMR) in the high coverage relative to the phase 3 panel. At an AF=5%, the improvement in INDEL imputation accuracy is still significant, ranging from 1.2-fold (EUR, SAS, AFR, EAS) to 1.4-fold (AMR), across the populations (Figure 5D). As expected, imputation performance for SNVs is comparable between the high coverage and phase 3 panels at AF > 5%. At AF < 5%, we observe slightly better performance with the high coverage panel across all 5 super-populations (e.g., SNVs at AF=0.2% are imputed with r^2^=0.54 using the high coverage panel vs. r^2^=0.45 with the phase 3 panel across EUR samples; Figure 5D).

## DISCUSSION

We present results from *high coverage* WGS of the expanded 1kGP cohort, consisting of 2,504 original samples as well as additional 698 related samples, completing 602 trios in the cohort. We called 111,048,944 SNVs, 14,435,076 INDELs, and 173,366 SVs across the 3,202 samples, using state-of-the-art methods. When compared to the low coverage phase 3 1kGP dataset published in 2015, the variant counts in the high coverage call set reflect an estimated average increase of 190,885 SNVs (1.05-fold), 268,182 INDELs (1.47-fold), and 5,835 (2.81-fold) SVs per genome, and a cohort-level increase of over 18.6 million SNV (1.24-fold), 9.8 million INDEL (4.05-fold), and ∼100 thousand SV loci (2.47-fold), across the original 2,504 unrelated samples. A *direct* comparison of the high coverage 1kGP SNV/INDEL dataset against the phase 3 set was impossible due to differences in genomic reference builds that were used for variant calling during generation of the two call sets. To enable a locus-level comparison, we lifted-over the phase 3 dataset from the GRCh37 to the GRCh38 reference, which was successful for 99.9% of phase 3 variants. Our goal was not to dissect all of the factors that likely influenced variant discovery in the high coverage and phase 3 datasets. Differences in sequencing platforms and read length (phase 3: Illumina HiSeq 2000 and HiSeq 2500, 76 bp or 101 bp paired-end reads; high coverage: Illumina NovaSeq 6000, 150 bp paired-end reads), library preparation (phase 3: PCR-based; high coverage: PCR-free), sequencing coverage (phase 3: mean depth 7.4X; high coverage: mean depth 34X), reference genome (phase 3: GRCh37; high coverage: GRCh38), alignment software (phase 3: BWA 0.5.9; high coverage: BWA-MEM 0.7.15), as well as in downstream bioinformatics pipeline most likely all contributed to various degrees to the differences in variant calls that we described here. Overall, despite these differences, we found high concordance between the high coverage and the phase 3 SNV/INDEL call set. Over 98% of small variants from the phase 3 call set were recalled in the high coverage dataset with AF correlation coefficient above 0.99 in the easy- and 0.93 in the difficult-to-sequence regions of the genome.

As expected, given that the phase 3 dataset identified nearly all common SNVs (MAF > 1%) in the population, the vast majority of the new SNVs identified here were in the rare MAF spectrum (≤ 1%). In terms of predicted functional SNVs, we called a comparable number of protein-coding variants relative to the phase 3 call set, which is in agreement with the expectation since the original call set included high coverage WES data in addition to low coverage WGS. We saw a more significant increase (∼1.2- to 1.4-fold) in the number of UTR and intronic SNVs, which typically exhibit lower coverage in WES. Consistent with the fact that high coverage sequencing and new variant callers bring greater improvements to INDEL as compared to SNV calling, we observed gains in INDEL counts across the entire MAF spectrum with gains in the rare end of the spectrum being the most pronounced. These gains in INDEL calls were apparent across all predicted functional categories, with cohort-level increases relative to the phase 3 call set ranging from 2.5-fold in start-loss category to 5-fold in the inframe insertion and stop-gain categories.

The SVs presented here provide a significant increase in discovery power over the phase 3 call set (9,655 vs. 3,431 SVs per genome) across all 1kGP populations. We further performed manual inspection of all large CNVs (> 50 kb, n=4,180), and benchmarked large inversions against Strand-seq (> 5 kb, n=250) to assess orthogonal support. Notably, an important advance from the SV discovery in this dataset is the updated prediction of functional alterations from SVs in each human genome. These analyses predicted that 162 protein coding genes were likely to be altered per genome across all SVs captured from short-read WGS. This updated result greatly exceeds estimates in the phase 3 call set (n=32 genes altered per genome). This prediction is comparable to prior estimates from SVs in short-read WGS from ∼15,000 individuals using blood-derived DNA in gnomAD (n=180 genes altered per genome (Collins et al., 2020)). If we consider the existing landscape of SVs from long-read WGS datasets in a subset of 34 of these samples, analyses from the HGSVC estimated an average of 24,653 SVs in each human genome, including 189 genes altered by SVs (Ebert et al., 2021). The data presented here, coupled with the availability of inheritance information from a large number of 602 complete trios in this dataset, and the independent long read WGS, Strand-seq, and optical mapping datasets on 34 of these samples from the HGSVC (Ebert et al., 2021), provides an unprecedented open access SV resource for methods development and genomic studies.

Inclusion of 602 trios in the expanded 1kGP cohort led to an order of magnitude greater accuracy of SNV/INDEL haplotype phasing due to both an increase in long-range haplotype sharing between related samples, and pedigree-based correction applied after statistical phasing to ensure consistency of phased haplotypes with the pedigree structure. Moreover, improvements in small variant calling, coupled with higher phasing accuracy of the high coverage panel, translated into significantly better imputation accuracy, especially for INDELs, across all of the 1kGP super-populations when the high coverage panel was used as the reference for imputation as compared to the phase 3 panel.

For more than a decade, the 1000 Genomes collection has been a key resource in the field of genomics. These datasets have produced scientific insights into population genetics and genome biology, as well as provided an openly sharable resource that has been widely used in methods development and testing as well as for technical validation. By generating high coverage sequencing data for the complete phase 3 set of 2,504 unrelated individuals and completing 602 trios with 698 additional samples, we have updated this critical resource with benchmarks and standards for a new generation of large-scale international whole genome sequencing initiatives. Our state of the art SNV, INDEL, and SV call sets, freely released, provide the most accurate and comprehensive catalog of variation compiled to date across this diverse genomic resource, particularly in rare SNVs and all classes of INDELs and SVs that were challenging to detect using earlier sequencing and analysis methods on low coverage data. We also present an improved phasing and imputation panel leveraging full sequence from trios that outperforms the existing imputation panel. Importantly, this panel is fully public and can be freely downloaded and used in combination with other panels and for use with any target dataset. Although many larger sequencing projects have now been conducted, the open nature of the 1000 Genomes samples will continue to make this a foundational resource for the community in the years to come.

## AUTHOR CONTRIBUTIONS

Writing of the manuscript and figure generation: M.B., U.S.E., X.Z., A.O.B. SNV/INDEL calling and analysis: U.S.E., M.B., R.M. SV calling: X.Z., H.B., H.A., A.A.R., A.C., W.E.C., The Human Genome Structural Variation Consortium. SV integration and analysis: X.Z., H.B. Haplotype phasing: M.B. Imputation evaluation: A.O.B. Production and quality control of the WGS data: L.W., A.R., U.S.E., M.B., K.N. Data coordination, data sharing, and user support: S.F., E.L., P.F., C.X. These authors jointly supervised this work: S.G., I.M.H., M.E.T, G.N., M.C.Z.

**The Human Genome Structural Variation Consortium:** Evan E. Eichler^1,2^, Jan O. Korbel^3^, Charles Lee^4,5^, Tobias Marschall^6^, Scott E. Devine^7^, William T. Harvey^1^, Weichen Zhou^8^, Ryan E. Mills^8,9^, Tobias Rausch^3^, Sushant Kumar^10,11^, Can Alkan^12,13^, Fereydoun Hormozdiari^14^, Zechen Chong^15^, Xiaofei Yang^4,16,17^, Jiadong Lin^18^, Mark B. Gerstein^10,11^, Ye Kai^17,18^, Qihui Zhu^4^, Feyza Yilmaz^4^, Chunlin Xiao^19^.

1. Department of Genome Sciences, University of Washington School of Medicine, Seattle, WA, USA.

2. Howard Hughes Medical Institute, University of Washington, Seattle, WA, USA.

3. European Molecular Biology Laboratory, Genome Biology Unit, Heidelberg, Germany.

4. The Jackson Laboratory for Genomic Medicine, Farmington, CT, USA.

5. Precision Medicine Center, The First Affiliated Hospital of Xi’an Jiaotong University, Xi’an, China.

6. Heinrich Heine University Düsseldorf, Medical Faculty, Institute for Medical Biometry and Bioinformatics, Düsseldorf, Germany.

7. Institute for Genome Sciences, University of Maryland School of Medicine, Baltimore, MD, USA.

8. Department of Computational Medicine & Bioinformatics, University of Michigan, Ann Arbor, MI, USA.

9. Department of Human Genetics, University of Michigan, Ann Arbor, MI, USA.

10. Department of Molecular Biophysics and Biochemistry, Yale University, New Haven, CT, USA.

11. Program in Computational Biology and Bioinformatics, Yale University, New Haven, CT, USA.

12. Department of Computer Engineering, Bilkent University, Ankara, Turkey.

13. Bilkent-Hacettepe Health Sciences and Technologies Program, Bilkent University, Ankara, Turkey.

14. Department of Biochemistry and Molecular Medicine, MIND Institute and Genome Center, University of California, Davis, CA, USA.

15. Department of Genetics and Informatics Institute, School of Medicine, University of Alabama at Birmingham, Birmingham, AL, USA.

16. Department of Computer Science and Technology, School of Electronic and Information Engineering, Xi’an Jiaotong University, Xi’an, China.

17. MOE Key Lab for Intelligent Networks & Networks Security, School of Electronics and Information Engineering, Xi’an Jiaotong University, Xi’an, China.

18. School of Automation Science and Engineering, Faculty of Electronic and Information Engineering, Xi’an Jiaotong University, Xi’an, China.

19. National Center for Biotechnology Information, National Library of Medicine, National Institutes of Health, Bethesda, MD, USA.

## METHODS

### RESOURCE AVAILABILITY

#### Lead contact

Requests for further information and resources should be directed to and will be fulfilled by the lead contact, Michael Zody.

#### Materials availability

This study did not generate new unique reagents.

#### Data and code availability

● FastQ files, CRAM alignment files, GVCFs, SNV/INDEL VCFs, SV VCF, haplotype-resolved SNV/INDEL VCFs, and sample metadata file listing pedigree and sex information for the 3,202 sequenced samples have been deposited in several public data repositories as described in https://www.internationalgenome.org/data-portal/data-collection/30x-grch38. The GRCh38 lifted-over version of the phase 3 1kGP SNV/INDEL call set, generated as part of this paper to facilitate comparative analysis, has been deposited on EBI FTP and is publicly available here: http://ftp.1000genomes.ebi.ac.uk/vol1/ftp/data_collections/1000G_2504_high_coverage/working/phase3_liftover_nygc_dir/
● This paper does not report original code.
● Any additional information required to reanalyze the data reported in this paper is available from the lead contact upon request.

### EXPERIMENTAL MODEL AND SUBJECT DETAILS

#### 1000 Genomes Project cohort

As part of this publication, we sequenced 3,202 lymphoblastoid cell line (LCL) samples from the 1kGP collection, including 1,598 males and 1,604 females. All cell lines were acquired from Coriell Institute for Medical Research. The 3,202 samples were drawn from 26 populations (listed in Table S1) across the following 5 continental ancestry groups: African (AFR, n=893), European (EUR, n=633), East Asian (EAS, n=601), South Asian (SAS, n=585), and American (AMR, n=490) (Figure 1A, Table S1). Among the 3,202 samples, there are 602 father-mother-child trios (including 2 trios that are part of a multi-generational family, and 10 trios that were split from 5 quads for the purpose of pedigree-based correction applied after haplotype phasing) and 6 parent-child duos. All reported relationships were confirmed in IBD analysis using KING v2.2.3 (Manichaikul et al., 2010).

### METHOD DETAILS

#### WGS library preparation and sequencing

DNA extracted from LCLs was ordered from the Coriell Institute for Medical Research for each of the 3,202 1kGP samples. Whole genome sequencing (WGS) libraries were prepared using the TruSeq DNA PCR-Free High Throughput Library Prep Kit in accordance with the manufacturer’s instructions. Briefly, 1ug of DNA was sheared using a Covaris LE220 sonicator (adaptive focused acoustics). DNA fragments underwent bead-based size selection and were subsequently end-repaired, adenylated, and ligated to Illumina sequencing adapters. Final libraries were evaluated using fluorescent-based assays including qPCR with the Universal KAPA Library Quantification Kit and Fragment Analyzer (Advanced Analytics) or BioAnalyzer (Agilent 2100). Libraries were sequenced on an Illumina NovaSeq 6000 system using 2 x 150bp cycles.

#### Quality control of sequence data

We ran a number of quality control (QC) tools to look for quality issues, sample swaps, and contamination issues. We ran FastQC (Andrews, 2019) v0.11.3 on the raw sequence data to assess yield and raw base qualities. We ran Picard (Broad Institute, 2019) v2.4.1 CollectMultipleMetrics and CollectWGSMetrics on the aligned BAM to collect alignment and insert size metrics. Picard CollectGcBiasMetrics was run to compute normalized coverage across multiple GC bins. Reads duplication metrics were quantified by running Picard MarkDuplicates on the BAM.

All the samples had at least 27X mean coverage across the genome (average per sample coverage: 34X, range: 27X-71X) and at least 91% of the bases at base quality score 30 or higher. The mean duplicate rate across the samples was 9% but there were 5 samples (HG00619, HG00982, HG02151, HG02573 and HG04039) that had a duplicate rate greater than 20. The median insert size per sample was 433 bp. Higher duplication rate is a known issue with Illumina’s patterned flow cell that uses exclusion amplification clustering method to increase data output, but this chemistry is very sensitive to library loading concentrations. Higher loading concentrations can lead to low throughput because of polyclonal clusters being formed in the nanowells of the patterned flow cell, whereas low concentration can lead to pad hopping which increases the duplication rate. VerifyBamID (Jun et al., 2012) was run in chip-free mode to estimate the likelihood of sample contamination. We use a cutoff of 2% to flag any sample for contamination and none of the samples reached the cutoff.

To make sure there were no sample mix-ups we ran genotype concordance against genotyping chip data. For that, we used the chip data that was released with phase 3. We did not find chip data for 15 samples in phase 3 so for those we ran Infinium CoreExome-24 v1.3 chip and performed genotype concordance. All the samples had > 97% genotype concordance.

#### SNV/INDEL discovery using GATK

Read alignment to the human reference genome GRCh38 using BWA-MEM v0.7.15 (Li, 2013), duplicate marking using Picard MarkDuplicates v2.4.1 (Broad Institute, 2019), and Base Quality Score Recalibration (BQSR) using GATK (McKenna et al., 2010) v3.5 BaseRecalibrator were performed according to the functional equivalence pipeline standard developed for the Centers for Common Disease Genomics project (Regier et al., 2018). SAM to BAM and BAM to CRAM file conversions were performed using Samtools v1.3.1 (Li et al., 2009). SNV and INDEL calling was performed using GATK (McKenna et al., 2010; Van der Auwera and O’Connor, 2020) v3.5, as described below. For variant discovery we used HaplotypeCaller in GVCF mode (Poplin et al., 2017) with sex-dependent ploidy settings on chromosome X and Y. Specifically, variant discovery on chromosome X was performed using diploid settings in females, diploid settings in PAR regions in males, and haploid settings in non-PAR regions in males. Variant discovery on chromosome Y was performed with haploid settings in males and was skipped entirely in females. We combined GVCFs in batches of ∼200 samples using GATK CombineGVCFs and jointly-genotyped all 3,202 samples with GenotypeGVCFs. We then used VariantRecalibrator to train the Variant Quality Score Recalibration (VQSR) model using “maxGaussians 8” and “maxGaussians 4” parameters for SNVs and INDELs, respectively. We applied the VQSR model to the joint call set using ApplyRecalibration with truth sensitivity levels of 99.8% for SNVs and 99.0% for INDELs.

#### Evaluation of small variant calls

BCFtools v1.9 (Li, 2011) was used to split multiallelic variants into multiple rows and left-normalize INDELs before counting variants at the cohort-level. Per sample variant metrics were collected using the GATK VariantEval tool (Van der Auwera and O’Connor, 2020). Mixed and complex variants and multi-nucleotide polymorphisms (MNPs) were not included in the breakdown of genome-level small variants. As part of QC, we estimated SNV density using the SNVDensity tool from VCFtools v0.1.12 (Danecek et al., 2011) in bins of 1000 bp across the callable genome, defined here as the GRCh38 reference excluding gaps (“N”s in the GRCh38 reference sequence). The mean SNV density across the callable genome (see Methods) is 39.46 per 1 kb of sequence. Chromosome 19 (43.21 SNVs per 1 kb) has the highest density overall across all chromosomes, whereas Chromosome X (30.16 SNVs per 1 kb) displays the lowest density across all chromosomes, followed by chromosome 1 (36.46 SNVs per 1 kb) among the autosomes which is in agreement with previous reports based on WGS data (Telenti et al., 2016).

We evaluated small variant calls separately in easy- and difficult-to-sequence regions of the genome, using stratification intervals defined by the GIAB (Krusche et al., 2019) and obtained from https://ftp-trace.ncbi.nlm.nih.gov/giab/ftp/release/genome-stratifications/v2.0/GRCh38/union/. Difficult regions include (i) tandem repeats and homopolymers longer than 6 bp (∼40% of difficult regions), (ii) segmental duplications (∼26% of difficult regions), (iii) low (< 25%) and high (> 65 %) GC content regions and “bad promoters” (∼39% of difficult regions), and (iv) regions with low mappability (∼39% of difficult regions) with some overlap between categories (https://ftp-trace.ncbi.nlm.nih.gov/giab/ftp/release/genome-stratifications/v2.0/GRCh38/union/v2.0-GRCh38-Union-README.txt). Any region in the GRCh38 reference that did not fall into a difficult region was classified as easy. The easy and difficult regions make up 80% and 20% of the reference genome, respectively.

FDR was estimated both genome-wide and in easy vs. difficult regions of the genome by comparing variant calls in sample NA12878 from the 3,202-sample high coverage call set to the GIAB NA12878 SNV/INDEL truth set v3.3.2 (Zook et al., 2019). The VCF files were compared using hap.py (v0.3.12; https://github.com/Illumina/hap.py) with the rtg-tools (v3.8.2) (Cleary et al., 2015) vcfeval comparison engine. All FDR calculations were restricted to the high confidence regions of the genome, as defined by the GIAB.

In addition to estimating FDR across all small variants and small variants in easy vs. difficult regions of the genome, we also estimated it among just the singletons. Due to the mixed nature of the expanded 1kGP cohort, which now includes both related and unrelated samples, the number of singletons (sites with AC=1 across the 3,202 samples) per sample varies depending on the sample’s relatedness status. This is because the sample’s relatedness status in the cohort (child, parent, unrelated) determines what the allele count of variants private to its family is across the sequenced cohort of samples (*i.e.* whether or not they are singletons in the cohort). In the high coverage call set, each child in a trio carries on average 19,795 inherited autosomal heterozygous variants that are shared only with one or both parents across all samples in the cohort. These variants can be further broken down into sites with AC=2 (mean=19,658), AC=3 (mean=135), and AC=4 (mean=1.75) within a trio. The mean number of variants private to a family per child in a trio closely matches the difference between the mean per-genome singleton count in children vs. unrelated samples, in agreement with the expectation (Figure S1E). Approximately half of these ∼20,000 sites are shared between the child and the mother and the other half between the child and the father, hence the mean singleton count in parents is halfway in between the mean singleton count in children and unrelated samples. Since sample NA12878 is a child in the expanded 1kGP cohort, we evaluated both its DNMs and variants private to the NA12878 trio to estimate FDR across singletons in the 3,202-sample jointly-genotyped high coverage call set. As additional validation, we estimated FDR among singletons in sample NA12878 from the independent jointly-genotyped high coverage call set of just the original 2,504 unrelated samples. Both of the FDR singleton analyses were restricted to the high confidence regions of the genome, as defined by either the GIAB v3.3.2 (Zook et al., 2019) or GIAB v4.2.1 (Wagner et al., 2021) truth sets.

Counts of assessed singletons in both of the FDR analysis:

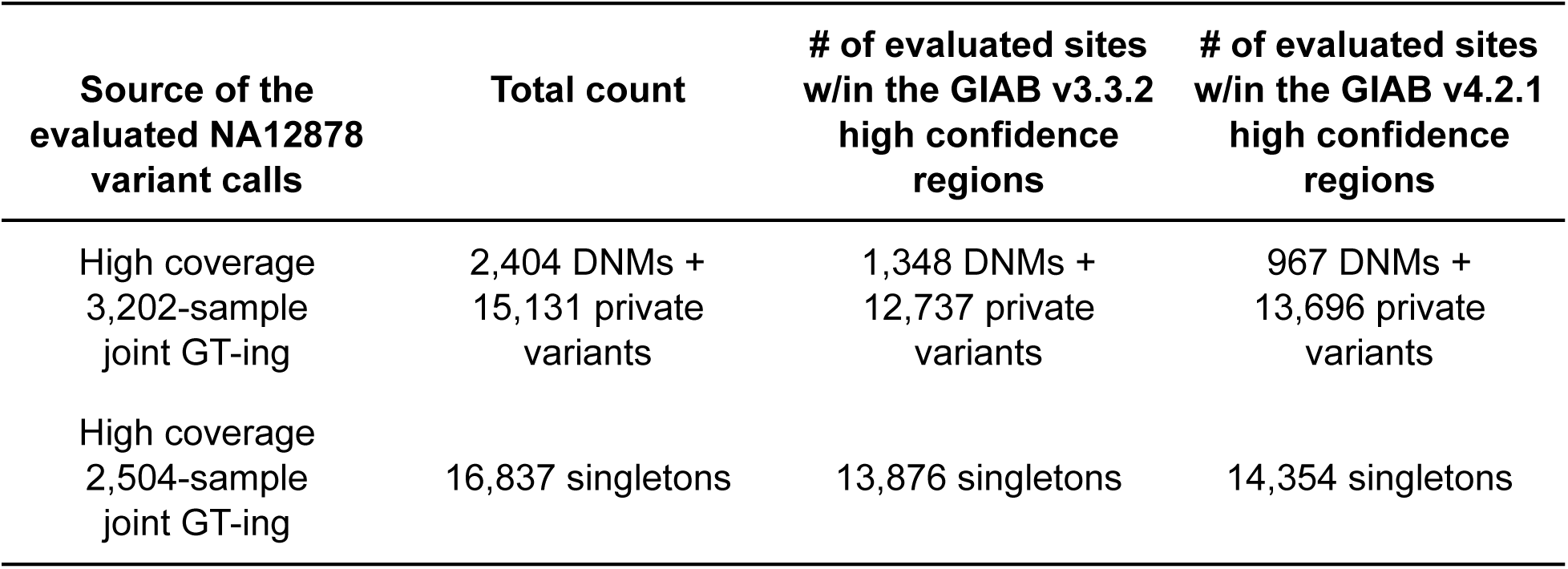

#### Functional consequence of small variants

We annotated small variant calls with predicted functional consequence using the Ensembl Variant Effect Predictor (VEP) v104 tool (McLaren et al., 2016). For each site, we chose one functional consequence per allele-gene combination (using “--pick_allele_gene” parameter) with default ordering of selection criteria. To avoid bias coming from families and to facilitate comparison to the phase 3 call set, cohort- and genome-level counts per predicted functional categories were reported based on the 2,504-sample jointly-genotyped high coverage call set which includes unrelated samples only (see Methods subsection below). Only variants that passed VQSR were considered in summary counts. No other filtering criteria were applied unless specifically noted.

#### Comparison of SNV/INDELs to the phase 3 set

To enable comparison of the high coverage against the phase 3 call set, we lifted-over the SNV/INDEL calls in the phase 3 call set from the GRCh37 to GRCh38 reference build using CrossMap v0.5.3 (Zhao et al., 2014). As input to the lift-over, we used the phase 3 VCFs available on the 1000 Genomes FTP, http://ftp.1000genomes.ebi.ac.uk/vol1/ftp/release/20130502/. Prior to the lift-over, we split multiallelic sites into separate rows. A small fraction of phase 3 loci (0.1%) failed the lift-over step due to the following reasons: 1) no hit found (unmapped GRCh37 variants); 2) loci mapping to multiple locations in the GRCh38 (multiple hits); 3) the reference allele matches the alternate allele after the lift-over (REF=ALT allele in the GRCh38). Additionally, we excluded variants that were lifted-over to a chromosome that was different from the original chromosome in GRCh37 (chromosome mismatch) or if the reference allele contained non-canonical nucleotide bases (non-canonical REF). Using this approach we were able to successfully lift-over 99.9% of phase 3 small variant loci (see table below). The resulting GRCh38 phase 3 call set that was used for the comparison was restricted to autosomes and contained 78,324,761 SNVs and 3,244,241 INDELs.

Table summarizing lift-over failures in the small variant phase 3 call set, consisting of 81,646,103 SNV/INDELs total:

| Failure reason | Count |
| --- | --- |
| REF=ALT allele in the GRCh38 | 27,889 |
| Unmapped GRCh37 variants | 15,856 |
| Multiple hits | 420 |
| Chromosome mismatch | 32,919 |
| Non-canonical REF | 17 |
| **Total lift-over failures** | **77,101 (0.1%)** |

We restricted the comparison of the high coverage vs. phase 3 calls to the 2,504 samples in common to the two cohorts. For that purpose, we generated an independent jointly-genotyped high coverage call set, including only the 2,504 original samples. Difference in FDR estimation between the 2,504- vs. 3,202-sample high coverage call set (0.1% vs. 0.3% for SNVs, respectively) is due to between-run variability caused by the non-deterministic nature of the VQSR step of the GATK SNV/INDEL calling pipeline (number of false positive SNVs across VQSR PASS sites: 4,098 vs. 9,227; number of false positive SNVs across all called sites: 22,807 vs. 22,994, in the 2,504- vs. 3,202-sample joint genotyping, respectively). The comparison of high coverage vs. phase 3 small variant call set was restricted to autosomes only. AF correlation across SNV and INDEL sites that are shared between the high coverage and the phase 3 call set was calculated using Pearson correlation coefficient obtained using the cor() function in R.

To compare the counts of small variants per functional consequence category between the high coverage and phase 3 call set, we annotated the GRCh38 lifted-over version of the phase 3 call set with the Ensembl VEP (the same way as described for the high coverage call set above), and computed ratios of cohort- and genome-level counts in the high coverage call set vs. phase 3 call set. To assess FDR across SNVs and INDELs in each functional category, we compared predicted functional SNVs and INDELs in the high coverage and phase 3 call sets to the GIAB NA12878 truth set v3.3.2 (Zook et al., 2019). The FDR calculation was restricted to the high confidence regions of the genome, as defined by the GIAB.

#### SV discovery using GATK-SV

GATK-SV involved an ensemble SV discovery and refinement pipeline for WGS data. The technical details of the method were previously described in Collins et al (Collins et al., 2020) for application to the genome aggregation database (gnomAD) for SV discovery, and further described in analyses from the HGSVC (Ebert et al., 2021). In this study, the same methods were applied to all 3,202 samples for SV discovery. In brief, SVs discovered by Manta, Wham, MELT, cn.MOPS and GATK-gCNV from Ebert et al. were integrated, genotyped across all samples, resolved for complex SVs, and annotated for variant class and functional impact. The FDR was previously assessed from analyses in quartet families, which yielded a 97% molecular validation rate for *de novo* SV predictions (Werling et al., 2018), as well as a 94% validation rate compared to long-read sequencing (Collins et al., 2020).

#### SV discovery using svtools

The svtools (Larson et al., 2019) method was previously described in (Abel et al., 2020) and applied for SV discovery across 17,795 genomes from the Centers for Common Disease Genomics (CCDG) program (Abel et al., 2020). The workflow combines per-sample variant discovery with lumpy (Layer et al., 2014) and manta (Chen et al., 2016) with resolution-aware cross-sample merging. The set of merged variants is then genotyped with svtyper (Chiang et al., 2015), followed by copy-number annotation with CNVnator (Abyzov et al., 2011) and reclassification of variants based on concordance of read-depth with breakpoint orientation. All parameter settings and versions are as implemented in the wdl-based workflow (https://github.com/hall-lab/sv-pipeline).

#### Large insertion discovery using Absinthe

On a per-sample basis, insertions with a minimum length of 100bp were discovered through *de novo* assembly of unmapped and discordant read pairs using Absinthe (Corvelo et al., 2021), and then genotyped using Paragraph (Chen et al., 2019), respecting sex-specific ploidies. Insertion calls from all 3,202 samples that were positively genotyped with a PASS filter flag were then clustered by genomic location and aligned using MAFFT (Katoh and Standley, 2013). For each locus, the most consensual allele was selected. Variants from the resulting merged call set were then re-genotyped on all 3,202 individuals. To produce the final call set only variants with 1) genotyping PASS filter rate ≥ 80%; 2) Mendelian Error Rate ≤ 5% for complete trio calls; and 3) HWE Chi-square test p-value > 1e-6 in at least one of the 5 super-populations were kept.

#### Integration of SV call sets

We conducted a series of analyses to benchmark SVs from each of the three methods described above, including their FDR as indicated by inheritance rates and support from orthogonal technologies, as well as their breakpoint precision estimated by the deviation of their SV breakpoints from long read assemblies in three genomes from analyses in the HGSVC (Chaisson et al., 2019). We also compared the three call sets to decide on the optimal integration strategy to maximize sensitivity and minimize FDR in the final ensemble call set (Figure S3, Table S5). Details of the comparison and integration strategies are described separately for insertions and all other variant classes below.

#### Integration of insertions

We compared the *de novo* rate of variant calls from each pipeline for insertions, yielding results of 4.1% for GATK-SV, 25.8% for svtools, and 2.4% for Absinthe. Given these results we restricted integration of insertions to GATK-SV and Absinthe. Each insertion pair was considered concordant if the insertion points were within 100 bp. The FDR of each insertion call set was estimated from three measurements: 1) *de novo* rate of SVs observed in the 602 trios; 2) proportion of SVs that were not validated by VaPoR (Zhao et al., 2017), an algorithm that evaluates SV quality by directly comparing raw PacBio reads against the reference genome, and 3) proportion of SVs that were not overlapped by SVs from PacBio assemblies in the same genome (Figure S3D-G). Precision of an insertion call was estimated by the distance of the insertion point to the closest PacBio insertion and the difference between the length of inserted sequence versus the length of the closest PacBio insertion calculated as an odds ratio. Both insertion call sets display less than 5% FDR based on inheritance and PacBio support, and the call sets were thus merged for all subsequent analyses (Figure S3D). Notably, as Absinthe showed higher precision than GATK-SV, as measured from both the coordinates of the insertion point and the length of inserted sequences (Figure S3H, I), we retained the Absinthe record for insertions that were shared by both methods.

#### Integration of SVs other than insertions

To consider a pair of SVs of the same variant class other than insertions as concordant, 50% reciprocal overlap was required for SVs larger than 5 kb and 10% reciprocal overlap was required for variants under 5 kb respectively. The FDR across variant calls was evaluated using the same measurements as described above. For deletions, duplications, and inversions, we observed low FDR (< 5%) among variants that were shared by GATK-SV and svtools, but significantly higher FDR in the subset that were uniquely discovered by either algorithm (Figure S3E-G). To restrict the final call set to high-quality variants, a machine learning model (lightGBM (Ke et al., 2017)) was trained on each SV class. Three samples that were previously analyzed in the HGSVC studies (HG00514, HG00733, NA19240) (Chaisson et al., 2019; Ebert et al., 2021) were selected to train the model. The truth data was defined by SVs that were uni-parentally inherited, shared by GATK-SV and svtools, supported by VaPoR, and overlapped by PacBio call sets. The false training subset was selected as SVs that appeared as *de novo* in offspring genomes, specifically discovered by either GATK-SV or svtools, not supported by VaPoR, and not overlapped by PacBio call sets. Multiple features were included in the model, including the sequencing depth of each SV, the depth of the 1 kb region around each SV, the count of aberrant pair ends (PE) within 150 bp of each SV, the count of split reads (SR) within 100 bp of each breakpoints, the size, allele fraction and genomic location (split into short repeats, segmental duplications, all remaining repeat masked regions, and the remaining unique sequences) of each SV, and the fraction of offspring harboring a *de novo* variant among trios in which the SV is observed. Each SV per genome was assigned a ‘boost score’ by the lightGBM model, and SVs with > 0.448 boost score were labeled as ‘PASS’ in the model (Figure S9M, N). This threshold was specifically selected to retain an estimated FDR < 5%. Call set specific SVs that failed the lightGBM model in less than 48% of all examined samples were included in the final integrated call set (Figure S3N).

To design strategies to merge SVs shared by GATK-SV and svtools, the precision of SV calls was evaluated by examining the distance between breakpoint coordinates of SVs to matched calls in the PacBio call set. Comparable breakpoint precision was observed for GATK-SV and svtools (Figure S3J-L). Thus, for SVs in each sample, the variant with the greatest number of split reads for each breakpoint was selected, or if equivalent then the variant with the higher boost score was retained, then for each locus the SV observed in the greatest number of samples was retained as final.

#### Inclusion of SVs exclusively from GATK-SV

Other minority SVs types, including mCNVs, CPX and CTX, were specifically detected by GATK-SV, so we performed in-depth manual inspection to ensure their quality before including them in the final integration call set. The depth profile across all 3,202 samples around each mCNV was plotted for manual review, and mCNVs that did not show clear stratification among samples were labeled as ‘Manual_LQ’ in the filter column even if they showed clear deviation from the normal copy number of 2. For CTX, the aberrantly aligned read-pairs across each breakpoint were manually examined, and variants that lacked sufficient support were labeled as ‘Manual_LQ’ in the final call set.

#### Comparison of SVs to the phase 3 call set

We compared the quality of SVs from the high-coverage WGS to the 1kGP phase 3 SV call set reported by Sudmant et al., 2015. Phase 3 SVs aligned against GRCh38 were obtained from the 1kGP ftp site: http://ftp.1000genomes.ebi.ac.uk/vol1/ftp/phase3/integrated_sv_map/supporting/GRCh38_positions/. It should be noted that 121 SVs failed liftover and were removed from the GRCh38 VCF, so a total of 68,697 SV sites were included in this comparison instead of 68,818 which was reported in Sudmant et al., 2015. When comparing SVs, we required 10% or higher reciprocal overlap for CNVs and INVs under 5 kb to be considered concordant, and 50% or higher reciprocal overlap for CNVs and INVs that are over 5 kb. We consider insertion pairs with insertion point within 100bp as concordant.

#### Haplotype phasing of small variants

To filter the SNV/INDEL call set for haplotype phasing, we first annotated the call set with HWE exact test p-values (Wigginton et al., 2005), stratified by super-population, using the BCFtools *fill-tags* plugin (Li, 2011). Next, we split multiallelic sites into separate rows and left-normalized representation of INDELs using BCFtools *norm* tool (Li, 2011). To ensure unique start position of all variant loci, required for phasing, we shifted positions of multiallelic sites by a minimum possible number of bp using an in-house script. The positions were shifted back to the original ones after phasing. Selection of SNVs and INDELs that passed VQSR, had GT missingness rate < 5%, passed HWE (*i.e.* had HWE exact test p-value > 1e-10 in at least one super-population), and had MAC ≥ 2 was done using BCFtools *(Li, 2011)*. Selection of variants with ME ≤ 5% was done using plink v1.90 (Purcell et al., 2007) after VCF to plink conversion (required to run phasing). For VCF to plink conversion we used plink v2.0 (Purcell et al., 2007). For haplotype phasing we used statistical phasing with pedigree-based correction, as implemented in SHAPEIT-duohmm v2.r904 (Delaneau et al., 2011; O’Connell et al., 2014). Phasing with SHAPEIT-duohmm was performed per chromosome using default settings, except for the window size parameter “-W” which was increased from 2Mb (default) to 5Mb to account for increased amounts of shared IBD due to pedigrees being present in the dataset (as recommended in the SHAPEIT manual). SHAPEIT-duohmm supports phasing of autosomal variants only. Therefore, to phase variants on chromosome X, we used statistical phasing as implemented in the Eagle v2.4.1 software (Loh et al., 2016). Phasing with Eagle was performed using default parameters. No shifting of positions for multiallelic sites was needed as Eagle supports phasing of variants with the same start site. Phasing accuracy evaluation was performed using the WhatsHap tool v0.18 (Martin et al., 2016). As a measure of phasing accuracy we used switch error rate (SER), which is defined as:

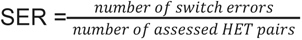

In all of the phasing evaluations, SER was computed across pairs of consecutive heterozygous sites in sample NA12878 (child in a trio in the expanded 3,202-sample cohort) relative to the Platinum Genome NA12878 gold standard truth set (Eberle et al., 2017).

#### Imputation performance evaluation

We performed imputation on 279 samples from 130 diverse populations using WGS data from the Simons Genome Diversity Project (SGDP) (Mallick et al., 2016). To create a pseudo-GWAS dataset, we extracted the genotypes at all sites included on the Illumina Infinium Omni2.5-8 v1.4 array. We performed quality control (QC) of the dataset using standard pre-imputation filters, removing sites which did not meet the following criteria: genotype call rate of ≥ 95%, MAF > 1%, and HWE p-value ≥ 1e-4. We used plink v1.9 (Purcell et al., 2007) for all QC steps, and analysis was restricted to the autosomes. We imputed the data passing quality control with the phase 3 and the high coverage reference panels, separately. The lifted-over GRCh38 phase 3 call set (described above) was used for the evaluations. We used SHAPEIT v2.r904 (Delaneau et al., 2011) to perform a strand check of the dataset and remove any problematic sites as determined by aligning with the respective panel. Pre-phasing was also performed using SHAPEIT and an input reference panel. We imputed the pre-phased data using IMPUTE v2.3.2 (Howie et al., 2009) software with default parameters. Following imputation, we concatenated the imputed intervals to create an autosome-wide imputed dataset. We evaluated imputation using 110 total samples with 22 samples from each of the five super-population ancestry groups (EUR, AFR, SAS, EAS, and AMR), the maximum number of samples available across all populations, and compared the imputed dosages with the WGS dosages stratified by MAF. For this evaluation, we converted the posterior genotype probabilities produced by IMPUTE v2.3.2 to dosages using QCTOOL v2.0.2 (https://www.well.ox.ac.uk/~gav/qctool_v2/), and the WGS genotypes to dosages using BCFtools v1.9 (Li, 2011). We then computed the correlation between the imputed dosages and those from the WGS data for all non-missing sites using squared Pearson correlation coefficient (r^2^; squared output of the cor() function in R). In the evaluation of the high coverage panel stratified by variant type and genomic regions (Figure 5C), the following numbers of variants were assessed: 45,140,382 SNVs and 2,229,248 INDELs (easy regions); 13,370,367 SNVs and 3,815,956 INDELs (difficult regions). To compare imputation accuracy between the phase 3 and the high coverage panels, we restricted the evaluations to sites that are shared between the two panels (49,758,888 SNVs and 2,298,294 INDELs; Figure 5D).

### QUANTIFICATION AND STATISTICAL ANALYSIS

Details of exact analyses, statistical tests, and tools can be found in the main text and Methods.

## ACKNOWLEDGEMENTS

The WGS data were generated at the New York Genome Center with funds provided by NHGRI Grants 3UM1HG008901-03S1 and 3UM1HG008901-04S1. A.C., W.E.C., and M.C.Z. were partially supported by the NHGRI grant UM1HG008901. S.F., E.L., and P.F. were partially supported by the Wellcome Trust (WT104947/Z/14/Z) and the European Molecular Biology Laboratory. Support for analyses by X.Z. and M.E.T. was provided by NIMH MH115957 and UM1HG008895. I.M.H, H.J.A. and A.A.R. were supported in part by the NHGRI grant UM1HG008853. C.X. was supported by the Intramural Research Program of the National Library of Medicine, National Institutes of Health. The HGSVC was supported in part by a grant from the National Institutes of Health (NIH) U24HG007497 (to C.L., E.E.E., J.O.K., T.M.). We thank Justin Zook from the Genome in a Bottle (GIAB) Consortium for his feedback and manual curation of a subset of singleton calls in the NA12878 sample across the technologies employed by the GIAB v4.2.1 truth set. All cell lines sequenced for this paper were obtained from Coriell Institute for Medical Research (NHGRI and NIGMS cell repositories). The following cell lines/DNA samples were obtained from the NIGMS Human Genetic Cell Repository at the Coriell Institute for Medical Research: [NA06984, NA06985, NA06986, NA06989, NA06991, NA06993, NA06994, NA06995, NA06997, NA07000, NA07014, NA07019, NA07022, NA07029, NA07031, NA07034, NA07037, NA07045, NA07048, NA07051, NA07055, NA07056, NA07340, NA07345, NA07346, NA07347, NA07348, NA07349, NA07357, NA07435, NA10830, NA10831, NA10835, NA10836, NA10837, NA10838, NA10839, NA10840, NA10842, NA10843, NA10845, NA10846, NA10847, NA10850, NA10851, NA10852, NA10853, NA10854, NA10855, NA10856, NA10857, NA10859, NA10860, NA10861, NA10863, NA10864, NA10865, NA11829, NA11830, NA11831, NA11832, NA11839, NA11840, NA11843, NA11881, NA11882, NA11891, NA11892, NA11893, NA11894, NA11917, NA11918, NA11919, NA11920, NA11930, NA11931, NA11932, NA11933, NA11992, NA11993, NA11994, NA11995, NA12003, NA12004, NA12005, NA12006, NA12043, NA12044, NA12045, NA12046, NA12056, NA12057, NA12058, NA12144, NA12145, NA12146, NA12154, NA12155, NA12156, NA12234, NA12239, NA12248, NA12249, NA12264, NA12272, NA12273, NA12274, NA12275, NA12282, NA12283, NA12286, NA12287, NA12329, NA12335, NA12336, NA12340, NA12341, NA12342, NA12343, NA12344, NA12347, NA12348, NA12375, NA12376, NA12383, NA12386, NA12399, NA12400, NA12413, NA12414, NA12485, NA12489, NA12546, NA12707, NA12708, NA12716, NA12717, NA12718, NA12739, NA12740, NA12748, NA12749, NA12750, NA12751, NA12752, NA12753, NA12760, NA12761, NA12762, NA12763, NA12766, NA12767, NA12775, NA12776, NA12777, NA12778, NA12801, NA12802, NA12812, NA12813, NA12814, NA12815, NA12817, NA12818, NA12827, NA12828, NA12829, NA12830, NA12832, NA12842, NA12843, NA12864, NA12865, NA12872, NA12873, NA12874, NA12875, NA12877, NA12878, NA12889, NA12890, NA12891, NA12892].

---

**Figure S1.**
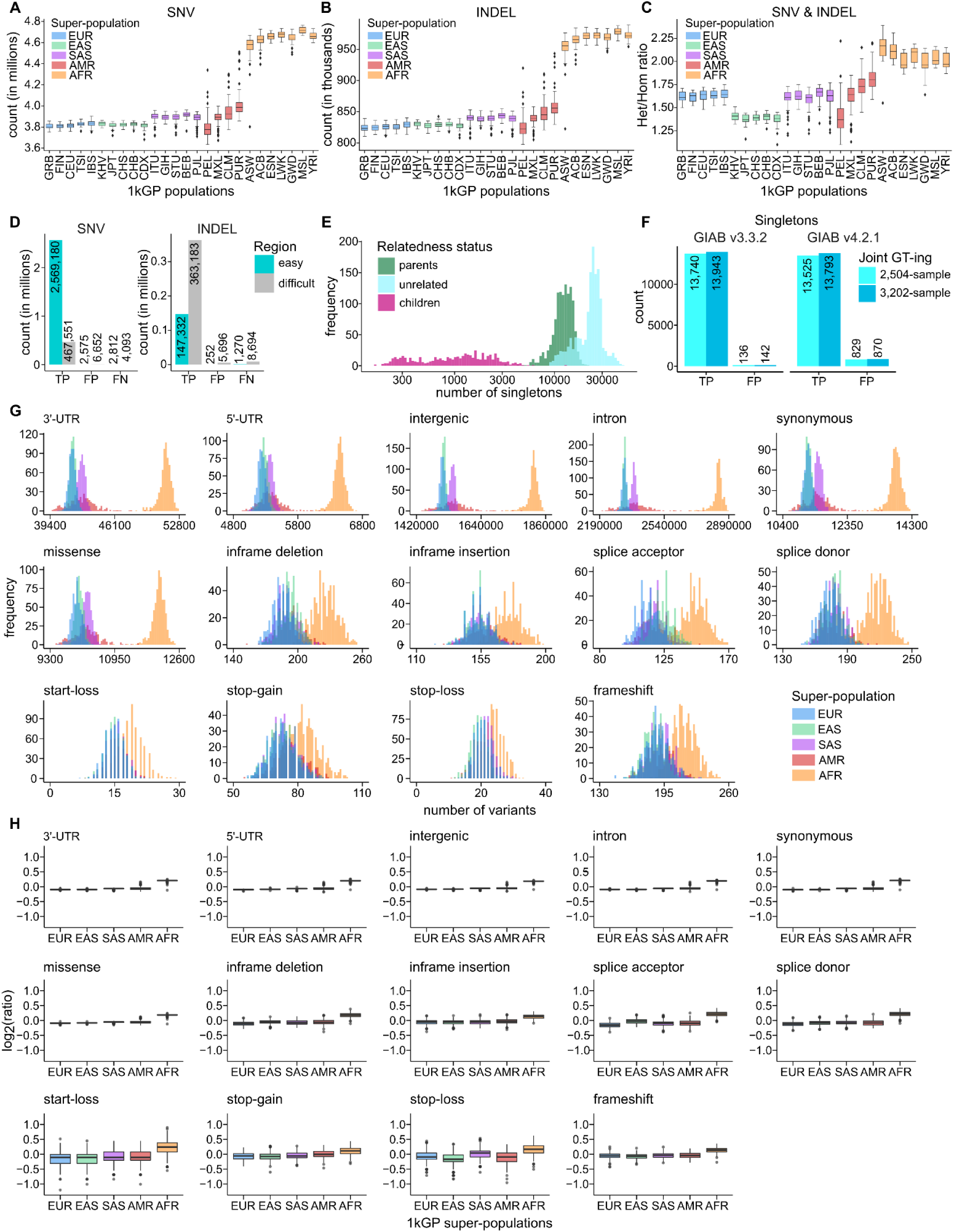
Evaluation of small variant calls, related to Figure 1. Genome-level counts of SNVs **(A)** and INDELs **(B)**, stratified by super-population. **(C)** Genome-level Het/Hom ratios across small variants, stratified by super-population. **(D)** Counts of true positive (TP), false positive (FP), and false negative (FN) SNV and INDEL calls in easy and difficult regions of the genome (GIAB v3.3.2 high confidence regions only). **(E)** Genome-level singleton (sites with AC=1 across 3,202 samples) counts across the 3,202 1kGP samples, stratified by relatedness status. **(F)** Counts of true positive (TP) and false positive (FP) singletons in NA12878 relative to either the GIAB v3.3.2 or GIAB v4.2.1 truth set (GIAB high confidence regions only). Due to the presence of NA12878’s parental samples in the expanded cohort, the analysis using the 3,202-sample 1kGP call set is based on both *de novos* and inherited variants private to the NA12878 trio. **(G)** Genome-level counts of predicted functional small variants, stratified by super-population. Reported counts are across the 2,504 unrelated samples only. **(H)** Distributions of log2(ratios) of genome-level counts from (G) normalized by the mean count across the 2,504 unrelated samples. Super-population ancestry labels: European (EUR), African (AFR), East Asian (EAS), South Asian (SAS), American (AMR). For descriptions of population labels please refer to Table S1.

**Figure S2.**
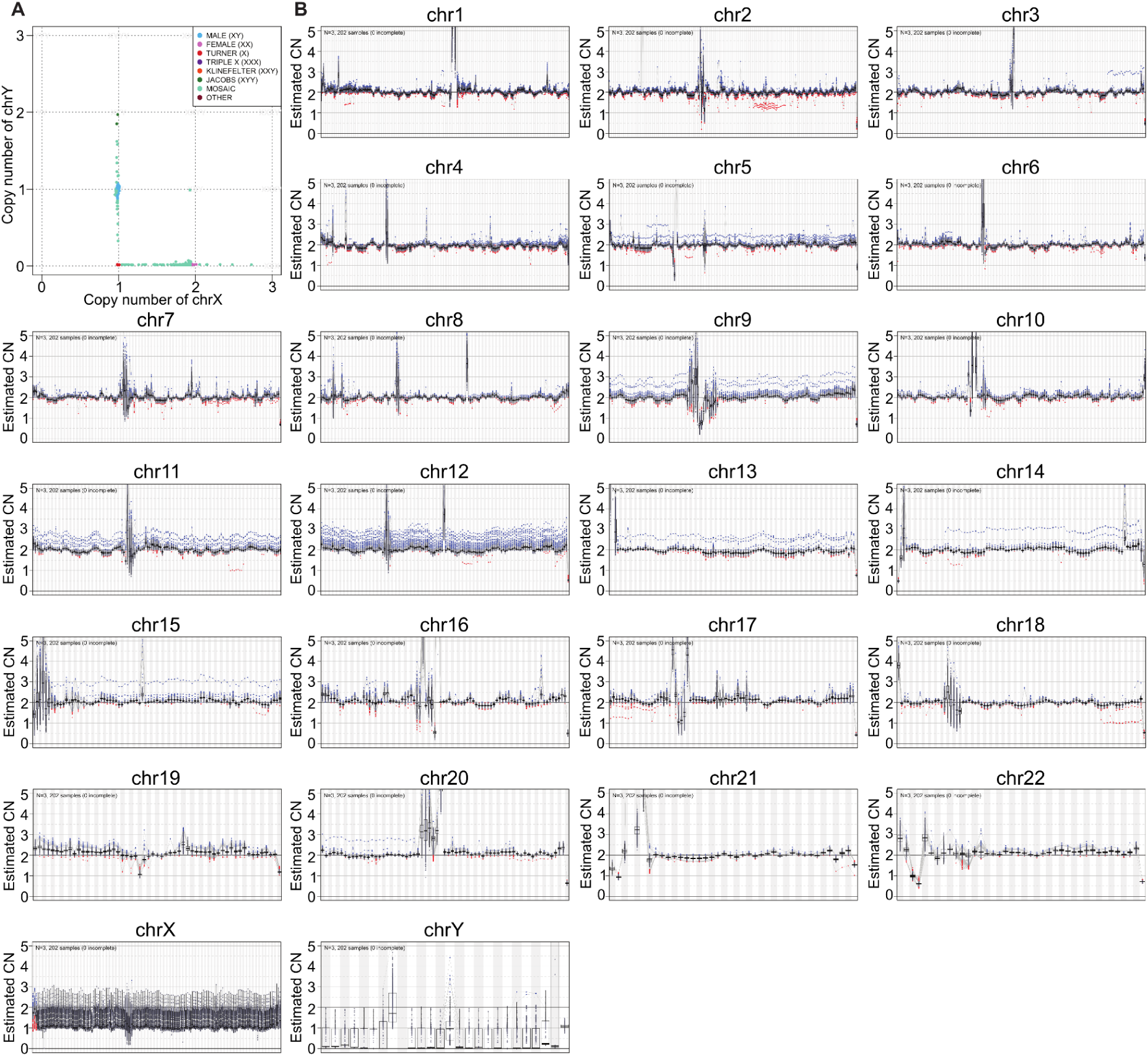
Ploidy of each chromosome across the 3,202 samples, related to Figure 1. **(A)** Ploidy of allosomes. **(B)** Copy number (CN) of each chromosome. Each dot represents a copy number of the 1Mbp bin in a sample. Blue dots are samples with copy gain and red dots represent copy loss.

**Figure S3.**
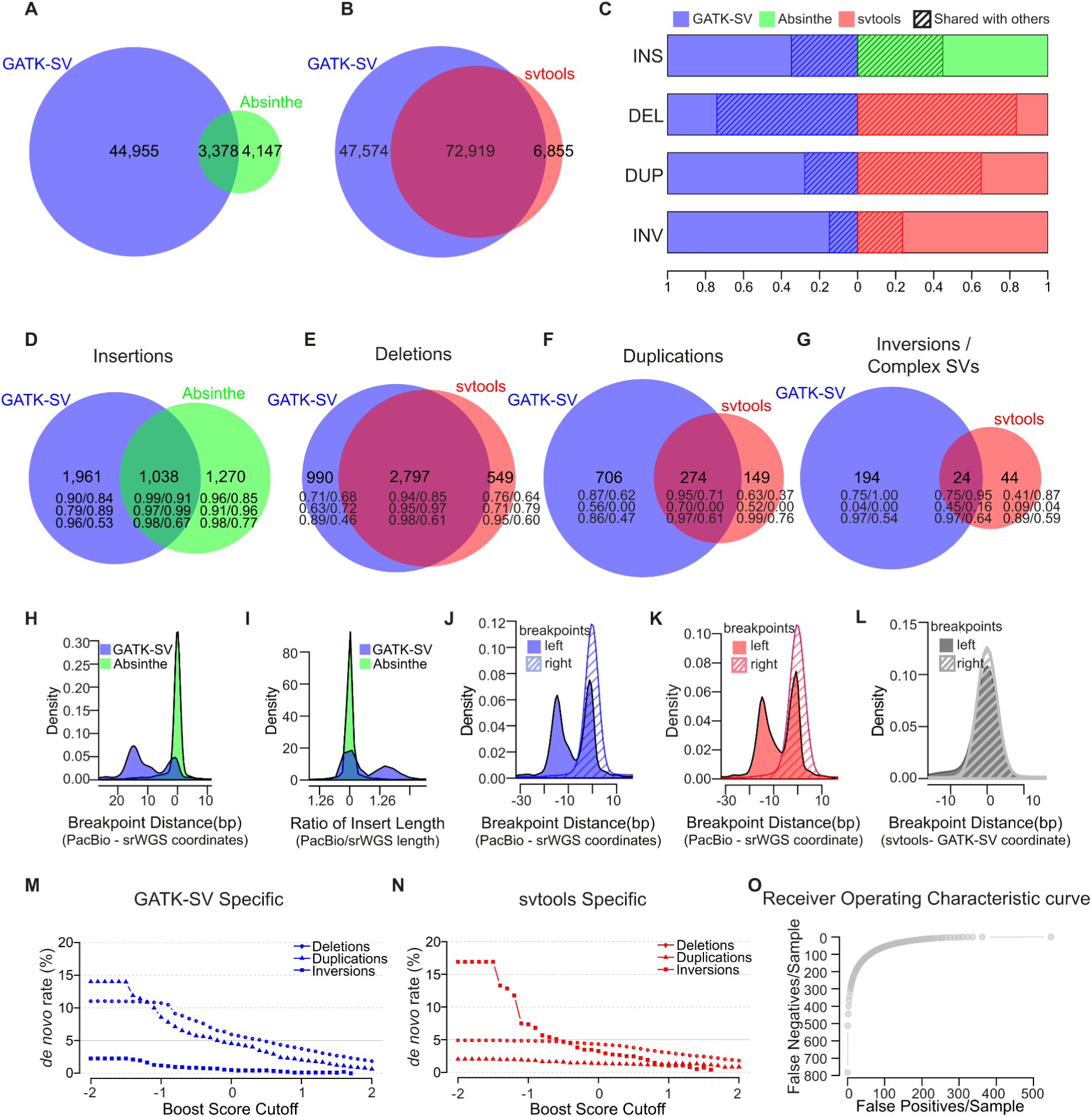
Benchmark of GATK-SV, svtools, and Absinthe, related to Figure 2. **(A)** Overlap of insertion sites between GATK-SV and Absinthe call sets. **(B)** Overlap of SV other than insertions between the GATK-SV and svtools call set. **(C)** Overlap of each SV type between GATK-SV, svtools, and Absinthe. **(D)** Overlap of insertions in each genome between GATK-SV and Absinthe. **(E-G)** Overlap of deletions (E), duplications (F), inversion and complex SVs (G) in each genome between GATK-SV and svtools. The integers in (D-G) represent count of SVs per sample, followed by proportion of SVs validated by VaPoR / proportion of SVs assessable by VaPoR in the second row, proportion of SVs supported by PacBio SVs in Ebert et al. 2021 / proportion of SVs supported by PacBio SVs in Chaisson *et al*. 2019 in the third row, and transmission rate /rate of bi-parentally inherited SVs in the fourth row. **(H-I)** Precision of the insertion breakpoint (H) and length (I) assessed against PacBio assemblies. **(J-K)** Precision of the SV breakpoints in GATK-SV (J) and svtools (K) call sets assessed against PacBio assemblies. **(L)** Breakpoint distance of SVs shared by GATK-SV and svtools. **(M-N)** *de novo* rate of SVs in GATK-SV (M) and svtools (N) call set when filtered at different boost score cutoffs. **(O)** False positives and false negatives in the GATK-SV and svtools call sets when filtered at different boost score cutoffs.

**Figure S4.**
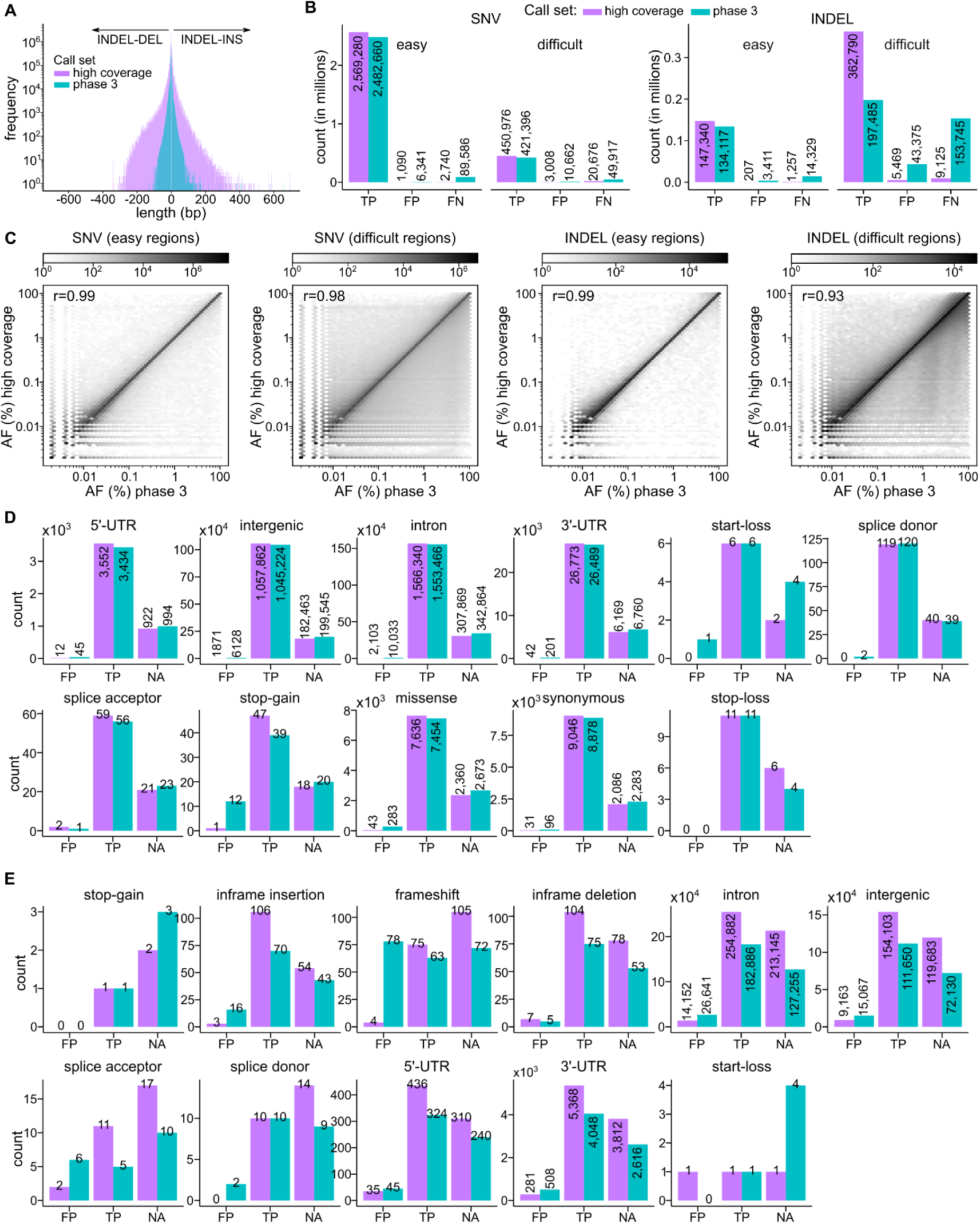
Comparison of small variant calls to the phase 3 call set, related to Figure 3. **(A)** Length of autosomal INDELs in the high coverage as compared to the phase 3 call sets. **(B)** Number of true positive (TP), false positive (FP), and false negative (FN) SNVs and INDELs in the high coverage vs. phase 3 call set, stratified by easy and difficult regions of the genome (GIAB v3.3.2 high confidence regions only). **(C)** Comparison of allele frequencies in the high coverage vs. the phase 3 call set across shared loci, stratified by variant type and easy vs. difficult regions of the genome. r: Pearson correlation coefficient. Number of false positive (FP), true positive (TP), and unassessed (NA; sites outside of the GIAB v3.3.2 high confidence regions of the genome) predicted functional SNVs **(D)** and INDELs **(E)** in sample NA12878, defined based on the comparison against the GIAB NA12878 truth set v3.3.2. See also Figure 3G, 3H (bottom row).

**Figure S5.**
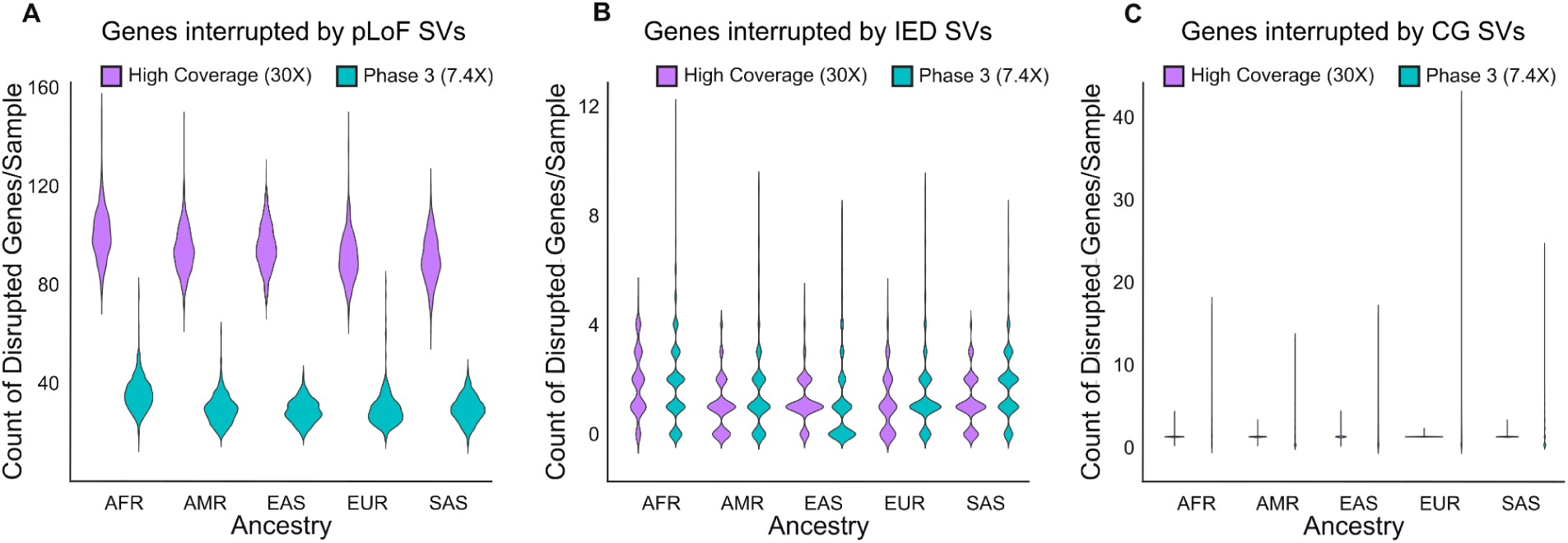
Comparison of gene interruptive SVs in the high-coverage ensemble versus low-coverage phase 3 1kGP call sets, related to Figure 4. **(A)** Count of genes interrupted as probable loss of function (pLoF), **(B)** intragenic exon duplications (IED), and **(C)** complete copy gain (CG) by SVs in the high-coverage ensemble call set and 1kGP phase 3 SV call set. Super-population ancestry labels: European (EUR), African (AFR), East Asian (EAS), South Asian (SAS), American (AMR).

**Figure S6.**
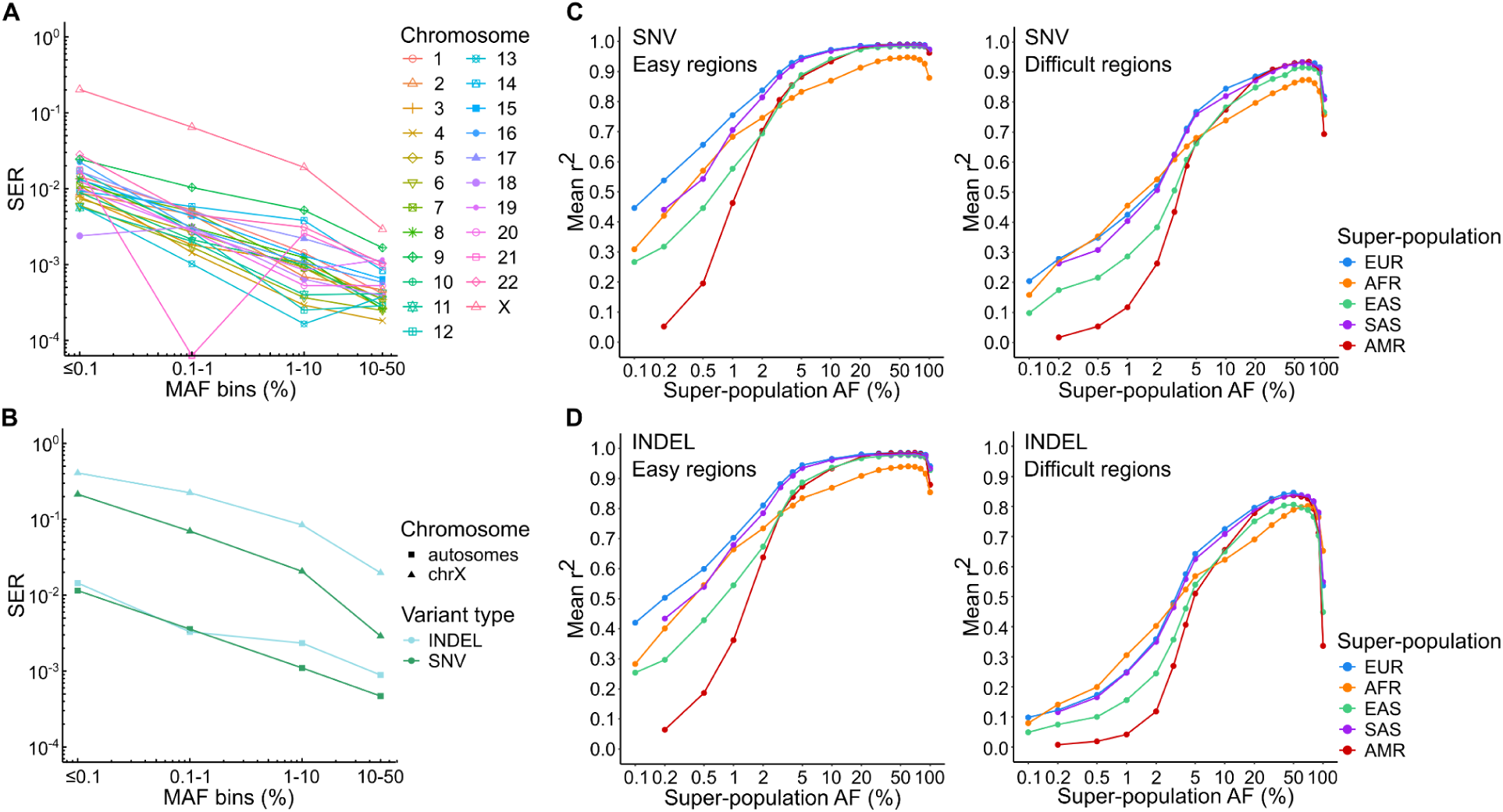
Evaluation of phasing and imputation performance of the high coverage 1kGP panel, related to Figure 5. SER: switch error rate stratified by **(A)** chromosome and **(B)** variant type. Chromosome X is shown separately in (B) as it was phased using a different strategy than autosomes (statistical phasing vs. statistical phasing with pedigree-based correction, respectively). Imputation accuracy of the high coverage panel stratified by super-population for SNVs **(C)** and INDELs **(D)** in easy and difficult regions of the genome. Imputation accuracy was estimated as described in Figure 5C.

**Table S1.** Sample counts in the expanded (3,202-sample) and original (2,504-sample) 1kGP cohort stratified by population, sex, and pedigree status, related to Figure 1. Thirteen samples are part of 2 trios (hence only 1,793 unique samples contribute to the 602 trios; not 1,806), either because they are part of a multi-generational family, *i.e*. are a child in one trio and a parent in another trio (HG00702, NA19685, NA19675), and/or because they are a part of a quad (5 quads were included in total) that was broken down into 2 trios when pedigree-based correction was applied following haplotype phasing (HG00656, HG00657, HG03642, HG03679, HG03943, HG03944, NA19660, NA19661, NA19678, NA19679). Super-population ancestry groups: European (EUR), African (AFR), East Asian (EAS), South Asian (SAS), American (AMR). Populations: African Caribbean in Barbados (ACB), People with African Ancestry in Southwest USA (ASW), Esan in Nigeria (ESN), Gambian in Western Division, Mandinka (GWD), Luhya in Webuye, Kenya (LWK), Mende in Sierra Leone (MSL), Yoruba in Ibadan, Nigeria (YRI), Colombians in Medellin, Colombia (CLM), People with Mexican Ancestry in Los Angeles, CA, USA (MXL), Peruvians in Lima, Peru (PEL), Puerto Ricans in Puerto Rico (PUR), Chinese Dai in Xishuangbanna, China (CDX), Han Chinese in Beijing, China (CHB), Han Chinese South, China (CHS), Japanese in Tokyo, Japan (JPT), Kinh in Ho Chi Minh City, Vietnam (KHV), Utah residents (CEPH) with Northern and Western European ancestry (CEU), Finnish in Finland (FIN), British from England and Scotland, UK (GBR), Iberian Populations in Spain (IBS), Toscani in Italia (TSI), Bengali in Bangladesh (BEB), Gujarati Indians in Houston, TX, USA (GIH), Indian Telugu in the UK (ITU), Punjabi in Lahore, Pakistan (PJL), Sri Lankan Tamil in the UK (STU).

| Population | Super-population | Sex<br>(1=male,<br>2=female) | # across<br>3,202<br>samples | # across<br>2,504<br>samples | # in trios | No. of trios |
| --- | --- | --- | --- | --- | --- | --- |
| ACB | AFR | 1 | 57 | 47 | 30 | 20 |
|  |  | 2 | 59 | 49 | 30 |  |
| ASW | AFR | 1 | 33 | 26 | 20 | 13 |
|  |  | 2 | 41 | 35 | 19 |  |
| ESN | AFR | 1 | 84 | 53 | 71 | 43 |
|  |  | 2 | 65 | 46 | 58 |  |
| GWD | AFR | 1 | 93 | 55 | 91 | 58 |
|  |  | 2 | 85 | 58 | 83 |  |
| LWK | AFR | 1 | 44 | 44 | 0 | 0 |
|  |  | 2 | 55 | 55 | 0 |  |
| MSL | AFR | 1 | 50 | 42 | 16 | 11 |
|  |  | 2 | 49 | 43 | 17 |  |
| YRI | AFR | 1 | 97 | 52 | 92 | 56 |
|  |  | 2 | 81 | 56 | 76 |  |
| CLM | AMR | 1 | 58 | 43 | 48 | 35 |
|  |  | 2 | 74 | 51 | 57 |  |
| MXL | AMR | 1 | 43 | 32 | 40 | 32 |
|  |  | 2 | 54 | 32 | 50 |  |
| PEL | AMR | 1 | 54 | 41 | 47 | 35 |
|  |  | 2 | 68 | 44 | 58 |  |
| PUR | AMR | 1 | 70 | 54 | 51 | 35 |
|  |  | 2 | 69 | 50 | 54 |  |
| CDX | EAS | 1 | 44 | 44 | 0 | 0 |
|  |  | 2 | 49 | 49 | 0 |  |
| CHB | EAS | 1 | 46 | 46 | 0 | 0 |
|  |  | 2 | 57 | 57 | 0 |  |
| CHS | EAS | 1 | 86 | 52 | 80 | 51 |
|  |  | 2 | 77 | 53 | 70 |  |
| JPT | EAS | 1 | 56 | 56 | 0 | 0 |
|  |  | 2 | 48 | 48 | 0 |  |
| KHV | EAS | 1 | 60 | 46 | 34 | 21 |
|  |  | 2 | 62 | 53 | 29 |  |
| CEU | EUR | 1 | 87 | 49 | 84 | 57 |
|  |  | 2 | 92 | 50 | 87 |  |
| FIN | EUR | 1 | 38 | 38 | 0 | 0 |
|  |  | 2 | 61 | 61 | 0 |  |
| GBR | EUR | 1 | 46 | 46 | 0 | 0 |
|  |  | 2 | 45 | 45 | 0 |  |
| IBS | EUR | 1 | 81 | 54 | 77 | 50 |
|  |  | 2 | 76 | 53 | 73 |  |
| TSI | EUR | 1 | 53 | 53 | 0 | 0 |
|  |  | 2 | 54 | 54 | 0 |  |
| BEB | SAS | 1 | 60 | 42 | 41 | 30 |
|  |  | 2 | 71 | 44 | 49 |  |
| GIH | SAS | 1 | 56 | 56 | 0 | 0 |
|  |  | 2 | 47 | 47 | 0 |  |
| ITU | SAS | 1 | 61 | 59 | 4 | 3 |
|  |  | 2 | 46 | 43 | 5 |  |
| PJL | SAS | 1 | 77 | 48 | 65 | 42 |
|  |  | 2 | 69 | 48 | 61 |  |
| STU | SAS | 1 | 65 | 55 | 16 | 10 |
|  |  | 2 | 49 | 47 | 10 |  |
| Total: |  |  | 3,202 | 2,504 | 1,793 | 602 |

**Table S2.** Mean SNV density per 1 kb of sequence in the 3,202-sample high coverage call set, related to Figure 1. Phased: SNV density in the phased high quality subset of SNV/INDEL calls; Genotyped: SNV density in the complete variant callset (based on VQSR PASS variants only).

| Chromosome | SNV Density per 1kb region |  |
| --- | --- | --- |
|  | Phased | Genotyped |
| chr1 | 21.88 | 36.46 |
| chr2 | 23.89 | 40.16 |
| chr3 | 23.89 | 40.11 |
| chr4 | 24.39 | 41.3 |
| chr5 | 23.76 | 39.91 |
| chr6 | 23.99 | 39.88 |
| chr7 | 24.61 | 41.1 |
| chr8 | 25.5 | 42.84 |
| chr9 | 21.76 | 36.66 |
| chr10 | 24.77 | 41.15 |
| chr11 | 24.11 | 40.61 |
| chr12 | 23.65 | 39.82 |
| chr13 | 24.22 | 41.04 |
| chr14 | 23.93 | 40.06 |
| chr15 | 23.52 | 39.21 |
| chr16 | 24.78 | 41.43 |
| chr17 | 23.39 | 39.02 |
| chr18 | 23.25 | 39.6 |
| chr19 | 26.62 | 43.21 |
| chr20 | 24.23 | 40.34 |
| chr21 | 22.84 | 37.77 |
| chr22 | 25.03 | 41.41 |
| chrX | 17.04 | 30.16 |

**Table S3.** Average sample-level count of small variants per functional consequence category stratified by super-population, related to Figure 1. Super-population ancestry labels: European (EUR), African (AFR), East Asian (EAS), South Asian (SAS), American (AMR).

| Filtering condition | Super-population | Functional consequence category |  |  |  |  |
| --- | --- | --- | --- | --- | --- | --- |
|  |  | stop-gain | splice donor & acceptor | frameshift | missense | synonymous |
| No filtering | All | 76 | 314 | 195 | 10,625 | 11,956 |
|  | AFR | 83 | 365 | 215 | 12,071 | 13,795 |
|  | EUR | 73 | 287 | 188 | 9,974 | 11,154 |
|  | EAS | 73 | 302 | 186 | 10,042 | 11,171 |
|  | SAS | 74 | 296 | 190 | 10,232 | 11,454 |
|  | AMR | 76 | 297 | 190 | 10,213 | 11,461 |
| MAF ≤ 1% | All | 11 | 18 | 14 | 754 | 569 |
|  | AFR | 15 | 30 | 22 | 1,215 | 1,044 |
|  | EUR | 9 | 12 | 10 | 540 | 353 |
|  | EAS | 10 | 14 | 12 | 594 | 398 |
|  | SAS | 10 | 14 | 12 | 648 | 453 |
|  | AMR | 10 | 13 | 11 | 567 | 387 |

**Table S4.** Count of SV sites at the cohort and sample level, related to Figure 2. SV types: DEL: deletion, DUP: duplication, mCNV: multiallelic copy number variant, INS: insertion, INV: inversion, BND: breakends, CPX: complex SV, CTX: inter-chromosomal translocation.

| SV TYPE | # SV sites across 3,202 samples |  |  | # SVs / sample |  |  |
| --- | --- | --- | --- | --- | --- | --- |
|  | GATK-SV | svtools | Absinthe | GATK-SV | svtools | Absinthe |
| INS | 48,333 | 75,283 | 7,183 | 3,019 | 1,761 | 2,270 |
| DEL | 89,445 | 65,184 | - | 3,783 | 3,417 | - |
| DUP | 26,353 | 10,594 | - | 990 | 459 | - |
| INV | 381 | 1,447 | - | 12 | 127 | - |
| BND | 82,218 | 26,152 | - | - | 2,188 | - |
| CPX | 3,624 | - | - | 216 | - | - |
| CTX | 16 | - | - | 1 | - | - |
| mCNV | 674 | - | - | 385 | - | - |
| ALL | 251,044 | 178,660 | 7,183 | 8,406 | 7,952 | 2,270 |

**Table S5.**
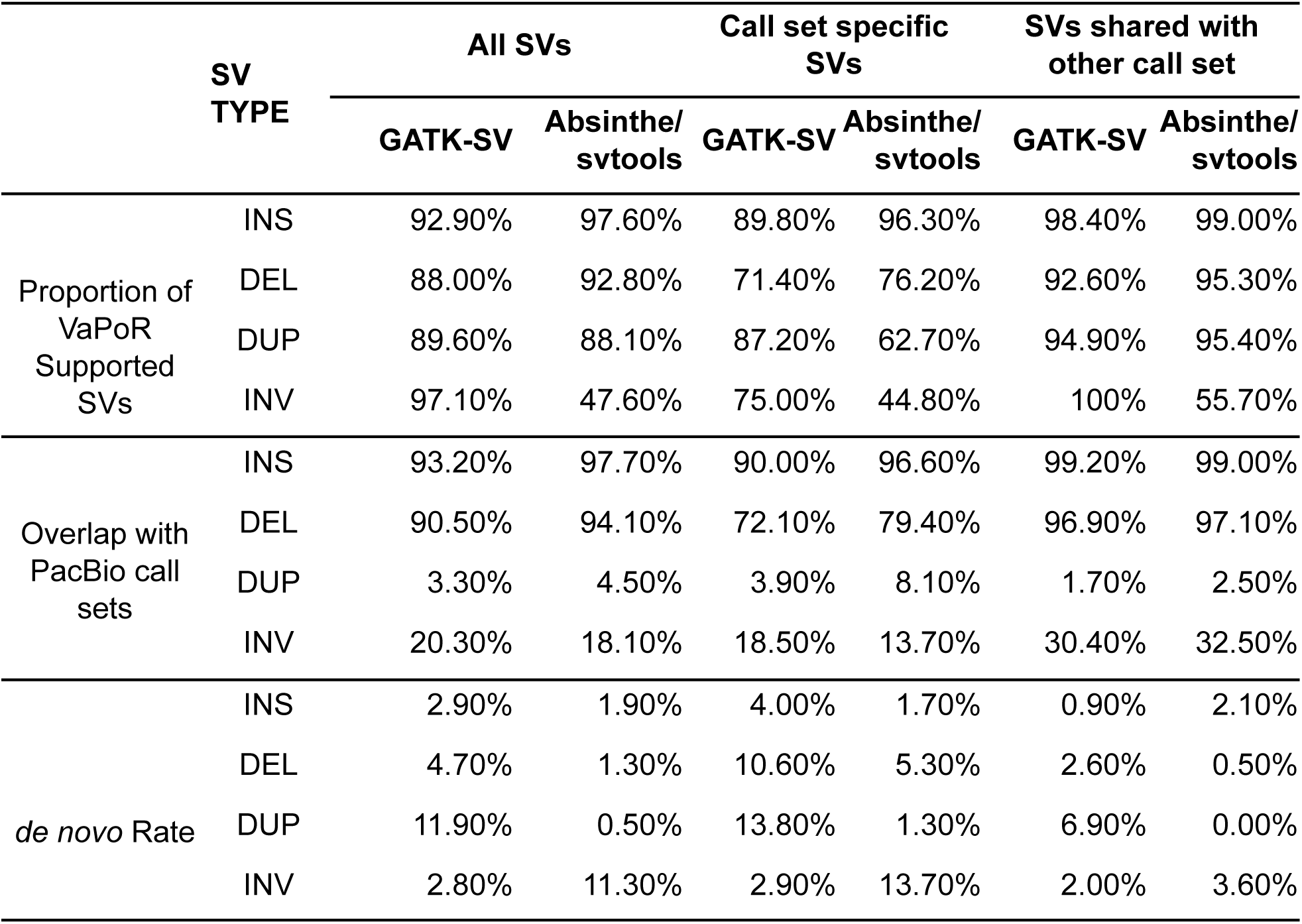
Quality of SVs evaluated by PacBio support and inheritance, related to Figure 2. SV types: INS: insertion, DEL: deletion, DUP: duplication, INV: inversion.

|  | SV<br>TYPE | All SVs |  | Call set specific<br>SVs |  | SVs shared with<br>other call set |  |
| --- | --- | --- | --- | --- | --- | --- | --- |
|  |  | GATK-SV | Absinthe/<br>svtools | GATK-SV | Absinthe/<br>svtools | GATK-SV | Absinthe/<br>svtools |
| Proportion of<br>VaPoR<br>Supported<br>SVs | INS | 92.90% | 97.60% | 89.80% | 96.30% | 98.40% | 99.00% |
|  | DEL | 88.00% | 92.80% | 71.40% | 76.20% | 92.60% | 95.30% |
|  | DUP | 89.60% | 88.10% | 87.20% | 62.70% | 94.90% | 95.40% |
|  | INV | 97.10% | 47.60% | 75.00% | 44.80% | 100% | 55.70% |
| Overlap with<br>PacBio call<br>sets | INS | 93.20% | 97.70% | 90.00% | 96.60% | 99.20% | 99.00% |
|  | DEL | 90.50% | 94.10% | 72.10% | 79.40% | 96.90% | 97.10% |
|  | DUP | 3.30% | 4.50% | 3.90% | 8.10% | 1.70% | 2.50% |
|  | INV | 20.30% | 18.10% | 18.50% | 13.70% | 30.40% | 32.50% |
| *de novo* Rate | INS | 2.90% | 1.90% | 4.00% | 1.70% | 0.90% | 2.10% |
|  | DEL | 4.70% | 1.30% | 10.60% | 5.30% | 2.60% | 0.50% |
|  | DUP | 11.90% | 0.50% | 13.80% | 1.30% | 6.90% | 0.00% |
|  | INV | 2.80% | 11.30% | 2.90% | 13.70% | 2.00% | 3.60% |

**Table S6.** Counts of small variants passing specified filtering criteria, related to Figure 5. Miss.: genotype-level missingness; HWE PASS: HWE exact test p-value > 1e-10 in at least one of the five 1kGP super-populations; ME: mendelian error rate across complete trios; MAC: minor allele count. MNP: multi-nucleotide polymorphism.

| **Filter** | **SNV** | **INDEL-DEL** | **INDEL-INS** | **MNP** |
| --- | --- | --- | --- | --- |
| VQSR PASS | 109,740,223 | 8,002,860 | 6,418,264 | 1,165,062 |
| Miss. < 5% | 108,637,798 | 9,588,168 | 7,044,553 | 1,132,357 |
| HWE PASS | 117,206,250 | 10,707,106 | 8,042,413 | 1,429,707 |
| ME ≤ 5% | 116,457,618 | 10,602,713 | 7,849,293 | 1,330,679 |
| MAC ≥ 2 | 71,444,181 | 7,671,955 | 6,302,585 | 1,188,242 |
| **All** | **61,411,215** | **5,400,097** | **4,554,384** | **699,618** |

## REFERENCES

1. Abel, H.J., Larson, D.E., Regier, A.A., Chiang, C., Das, I., Kanchi, K.L., Layer, R.M., Neale, B.M., Salerno, W.J., Reeves, C., et al. (2020). Mapping and characterization of structural variation in 17,795 human genomes. Nature 583, 83–89.

2. Abyzov, A., Urban, A.E., Snyder, M., and Gerstein, M. (2011). CNVnator: an approach to discover, genotype, and characterize typical and atypical CNVs from family and population genome sequencing. Genome Res. 21, 974–984.

3. Almeida, R., Ricaño-Ponce, I., Kumar, V., Deelen, P., Szperl, A., Trynka, G., Gutierrez-Achury, J., Kanterakis, A., Westra, H.-J., Franke, L., et al. (2014). Fine mapping of the celiac disease-associated LPP locus reveals a potential functional variant. Hum. Mol. Genet. 23, 2481–2489.

4. Andrews, S. (2019). FastQC (https://www.bioinformatics.babraham.ac.uk/projects/fastqc/).

5. Broad Institute (2019). Picard Toolkit, Github repository (http://broadinstitute.github.io/picard/).

6. Campbell, M.C., and Tishkoff, S.A. (2008). African genetic diversity: implications for human demographic history, modern human origins, and complex disease mapping. Annu. Rev. Genomics Hum. Genet. 9, 403–433.

7. Chaisson, M.J.P., Sanders, A.D., Zhao, X., Malhotra, A., Porubsky, D., Rausch, T., Gardner, E.J., Rodriguez, O.L., Guo, L., Collins, R.L., et al. (2019). Multi-platform discovery of haplotype-resolved structural variation in human genomes. Nat. Commun. 10, 1784.

8. Chen, S., Krusche, P., Dolzhenko, E., Sherman, R.M., Petrovski, R., Schlesinger, F., Kirsche, M., Bentley, D.R., Schatz, M.C., Sedlazeck, F.J., et al. (2019). Paragraph: a graph-based structural variant genotyper for short-read sequence data. Genome Biol. 20, 291.

9. Chen, X., Schulz-Trieglaff, O., Shaw, R., Barnes, B., Schlesinger, F., Källberg, M., Cox, A.J., Kruglyak, S., and Saunders, C.T. (2016). Manta: rapid detection of structural variants and indels for germline and cancer sequencing applications. Bioinformatics 32, 1220–1222.

10. Chiang, C., Layer, R.M., Faust, G.G., Lindberg, M.R., Rose, D.B., Garrison, E.P., Marth, G.T., Quinlan, A.R., and Hall, I.M. (2015). SpeedSeq: ultra-fast personal genome analysis and interpretation. Nat. Methods 12, 966–968.

11. Cleary, J.G., Braithwaite, R., Gaastra, K., Hilbush, B.S., Inglis, S., Irvine, S.A., Jackson, A., Littin, R., Rathod, M., Ware, D., et al. (2015). Comparing Variant Call Files for Performance Benchmarking of Next-Generation Sequencing Variant Calling Pipelines. bioRxiv 023754.

12. Collins, R.L., Brand, H., Karczewski, K.J., Zhao, X., Alföldi, J., Francioli, L.C., Khera, A.V., Lowther, C., Gauthier, L.D., Wang, H., et al. (2020). A structural variation reference for medical and population genetics. Nature 581, 444–451.

13. Corvelo, A., Clarke, W.E., Zody, M.C. (2021). Absinthe, Github repository (github.com/nygenome/absinthe).

14. Danecek, P., Auton, A., Abecasis, G., Albers, C.A., Banks, E., DePristo, M.A., Handsaker, R.E., Lunter, G., Marth, G.T., Sherry, S.T., et al. (2011). The variant call format and VCFtools. Bioinformatics 27, 2156–2158.

15. Delaneau, O., Marchini, J., and Zagury, J.-F. (2011). A linear complexity phasing method for thousands of genomes. Nat. Methods 9, 179–181.

16. Eberle, M.A., Fritzilas, E., Krusche, P., Källberg, M., Moore, B.L., Bekritsky, M.A., Iqbal, Z., Chuang, H.-Y., Humphray, S.J., Halpern, A.L., et al. (2017). A reference data set of 5.4 million phased human variants validated by genetic inheritance from sequencing a three-generation 17-member pedigree. Genome Res. 27, 157–164.

17. Ebert, P., Audano, P.A., Zhu, Q., Rodriguez-Martin, B., Porubsky, D., Bonder, M.J., Sulovari, A., Ebler, J., Zhou, W., Serra Mari, R., et al. (2021). Haplotype-resolved diverse human genomes and integrated analysis of structural variation. Science 372, eabf7117.

18. Hara, K., Fujita, H., Johnson, T.A., Yamauchi, T., Yasuda, K., Horikoshi, M., Peng, C., Hu, C., Ma, R.C.W., Imamura, M., et al. (2014). Genome-wide association study identifies three novel loci for type 2 diabetes. Hum. Mol. Genet. 23, 239–246.

19. Horikoshi, M., Mӓgi, R., van de Bunt, M., Surakka, I., Sarin, A.-P., Mahajan, A., Marullo, L., Thorleifsson, G., Hӓgg, S., Hottenga, J.-J., et al. (2015). Discovery and Fine-Mapping of Glycaemic and Obesity-Related Trait Loci Using High-Density Imputation. PLoS Genet. 11, e1005230.

20. Howie, B.N., Donnelly, P., and Marchini, J. (2009). A flexible and accurate genotype imputation method for the next generation of genome-wide association studies. PLoS Genet. 5, e1000529.

21. Huang, J., Chen, J., Esparza, J., Ding, J., Elder, J.T., Abecasis, G.R., Lee, Y.-A., Mark Lathrop, G., Moffatt, M.F., Cookson, W.O.C., et al. (2015). eQTL mapping identifies insertion- and deletion-specific eQTLs in multiple tissues. Nat. Commun. 6, 6821.

22. Illumina Inc. (2019). Polaris, Github repository (https://github.com/Illumina/Polaris/tree/master/cohorts/1000_genomes).

23. Jónsson, H., Sulem, P., Kehr, B., Kristmundsdottir, S., Zink, F., Hjartarson, E., Hardarson, M.T., Hjorleifsson, K.E., Eggertsson, H.P., Gudjonsson, S.A., et al. (2017). Parental influence on human germline de novo mutations in 1,548 trios from Iceland. Nature 549, 519–522.

24. Jun, G., Flickinger, M., Hetrick, K.N., Romm, J.M., Doheny, K.F., Abecasis, G.R., Boehnke, M., and Kang, H.M. (2012). Detecting and Estimating Contamination of Human DNA Samples in Sequencing and Array-Based Genotype Data. The American Journal of Human Genetics 91, 839–848.

25. Karczewski, K.J., Francioli, L.C., Tiao, G., Cummings, B.B., Alföldi, J., Wang, Q., Collins, R.L., Laricchia, K.M., Ganna, A., Birnbaum, D.P., et al. (2020). The mutational constraint spectrum quantified from variation in 141,456 humans. Nature 581, 434–443.

26. Katoh, K., and Standley, D.M. (2013). MAFFT multiple sequence alignment software version 7: improvements in performance and usability. Mol. Biol. Evol. 30, 772–780.

27. Ke, G., Meng, Q., Finley, T., Wang, T., Chen, W., Ma, W., Ye, Q., and Liu, T.-Y. (2017). LightGBM: A Highly Efficient Gradient Boosting Decision Tree. In Advances in Neural Information Processing Systems 30, I. Guyon, U.V. Luxburg, S. Bengio, H. Wallach, R. Fergus, S. Vishwanathan, and R. Garnett, eds. (Curran Associates, Inc.), pp. 3146–3154.

28. Khurana, E., Fu, Y., Colonna, V., Mu, X.J., Kang, H.M., Lappalainen, T., Sboner, A., Lochovsky, L., Chen, J., Harmanci, A., et al. (2013). Integrative annotation of variants from 1092 humans: application to cancer genomics. Science 342, 1235587.

29. Kircher, M., Witten, D.M., Jain, P., O’Roak, B.J., Cooper, G.M., and Shendure, J. (2014). A general framework for estimating the relative pathogenicity of human genetic variants. Nat. Genet. 46, 310–315.

30. Kong, A., Frigge, M.L., Masson, G., Besenbacher, S., Sulem, P., Magnusson, G., Gudjonsson, S.A., Sigurdsson, A., Jonasdottir, A., Jonasdottir, A., et al. (2012). Rate of de novo mutations and the importance of father’s age to disease risk. Nature 488, 471–475.

31. Krusche, P., Trigg, L., Boutros, P.C., Mason, C.E., De La Vega, F.M., Moore, B.L., Gonzalez-Porta, M., Eberle, M.A., Tezak, Z., Lababidi, S., et al. (2019). Best practices for benchmarking germline small-variant calls in human genomes. Nat. Biotechnol. 37, 555–560.

32. Lappalainen, T., Sammeth, M., Friedländer, M.R., ’t Hoen, P.A.C., Monlong, J., Rivas, M.A., Gonzàlez-Porta, M., Kurbatova, N., Griebel, T., Ferreira, P.G., et al. (2013). Transcriptome and genome sequencing uncovers functional variation in humans. Nature 501, 506–511.

33. Larson, D.E., Abel, H.J., Chiang, C., Badve, A., Das, I., Eldred, J.M., Layer, R.M., and Hall, I.M. (2019). svtools: population-scale analysis of structural variation. Bioinformatics 35, 4782–4787.

34. Layer, R.M., Chiang, C., Quinlan, A.R., and Hall, I.M. (2014). LUMPY: a probabilistic framework for structural variant discovery. Genome Biol. 15, R84.

35. Li, H. (2011). A statistical framework for SNP calling, mutation discovery, association mapping and population genetical parameter estimation from sequencing data. Bioinformatics 27, 2987–2993.

36. Li, H. (2013). Aligning sequence reads, clone sequences and assembly contigs with BWA-MEM. arXiv:1303.3997v1 [q-bio.GN].

37. Li, H., Handsaker, B., Wysoker, A., Fennell, T., Ruan, J., Homer, N., Marth, G., Abecasis, G., Durbin, R., and 1000 Genome Project Data Processing Subgroup (2009). The Sequence Alignment/Map format and SAMtools. Bioinformatics 25, 2078–2079.

38. Loh, P.-R., Danecek, P., Palamara, P.F., Fuchsberger, C., A Reshef, Y., K Finucane, H., Schoenherr, S., Forer, L., McCarthy, S., Abecasis, G.R., et al. (2016). Reference-based phasing using the Haplotype Reference Consortium panel. Nat. Genet. 48, 1443–1448.

39. Mallick, S., Li, H., Lipson, M., Mathieson, I., Gymrek, M., Racimo, F., Zhao, M., Chennagiri, N., Nordenfelt, S., Tandon, A., et al. (2016). The Simons Genome Diversity Project: 300 genomes from 142 diverse populations. Nature 538, 201–206.

40. Manichaikul, A., Mychaleckyj, J.C., Rich, S.S., Daly, K., Sale, M., and Chen, W.-M. (2010). Robust relationship inference in genome-wide association studies. Bioinformatics 26, 2867–2873.

41. Martin, M., Patterson, M., Garg, S., Fischer, S.O., Pisanti, N., Klau, G.W., Schöenhuth, A., and Marschall, T. (2016). WhatsHap: fast and accurate read-based phasing. bioRxiv 085050.

42. McKenna, A., Hanna, M., Banks, E., Sivachenko, A., Cibulskis, K., Kernytsky, A., Garimella, K., Altshuler, D., Gabriel, S., Daly, M., et al. (2010). The Genome Analysis Toolkit: a MapReduce framework for analyzing next-generation DNA sequencing data. Genome Res. 20, 1297–1303.

43. McLaren, W., Gil, L., Hunt, S.E., Riat, H.S., Ritchie, G.R.S., Thormann, A., Flicek, P., and Cunningham, F. (2016). The Ensembl Variant Effect Predictor. Genome Biol. 17, 122.

44. Montgomery, S.B., Goode, D.L., Kvikstad, E., Albers, C.A., Zhang, Z.D., Mu, X.J., Ananda, G., Howie, B., Karczewski, K.J., Smith, K.S., et al. (2013). The origin, evolution, and functional impact of short insertion-deletion variants identified in 179 human genomes. Genome Research 23, 749–761.

45. Ng, J.K., Vats, P., Fritz-Waters, E., Sarkar, S., Sams, E.I., Padhi, E.M., Payne, Z.L., Leonard, S., West, M.A., Prince, C., et al. (2021). de novo variant calling identifies cancer mutation profiles in the 1000 Genomes Project. bioRxiv 445979.

46. Nikpay, M., Goel, A., Won, H.-H., Hall, L.M., Willenborg, C., Kanoni, S., Saleheen, D., Kyriakou, T., Nelson, C.P., Hopewell, J.C., et al. (2015). A comprehensive 1000 Genomes–based genome-wide association meta-analysis of coronary artery disease. Nat. Genet. 47, 1121–1130.

47. O’Connell, J., Gurdasani, D., Delaneau, O., Pirastu, N., Ulivi, S., Cocca, M., Traglia, M., Huang, J., Huffman, J.E., Rudan, I., et al. (2014). A general approach for haplotype phasing across the full spectrum of relatedness. PLoS Genet. 10, e1004234.

48. Poplin, R., Ruano-Rubio, V., DePristo, M.A., Fennell, T.J., Carneiro, M.O., Van der Auwera, G.A., Kling, D.E., Gauthier, L.D., Levy-Moonshine, A., Roazen, D., et al. (2017). Scaling accurate genetic variant discovery to tens of thousands of samples. bioRxiv Doi: 10.1101/201178 201178.

49. Purcell, S., Neale, B., Todd-Brown, K., Thomas, L., Ferreira, M.A.R., Bender, D., Maller, J., Sklar, P., de Bakker, P.I.W., Daly, M.J., et al. (2007). PLINK: a tool set for whole-genome association and population-based linkage analyses. Am. J. Hum. Genet. 81, 559–575.

50. Regier, A.A., Farjoun, Y., Larson, D.E., Krasheninina, O., Kang, H.M., Howrigan, D.P., Chen, B.-J., Kher, M., Banks, E., Ames, D.C., et al. (2018). Functional equivalence of genome sequencing analysis pipelines enables harmonized variant calling across human genetics projects. Nat. Commun. 9, 4038.

51. Ritchie, G.R.S., Dunham, I., Zeggini, E., and Flicek, P. (2014). Functional annotation of noncoding sequence variants. Nat. Methods 11, 294–296.

52. Sherry, S.T., Ward, M., and Sirotkin, K. (1999). dbSNP—Database for Single Nucleotide Polymorphisms and Other Classes of Minor Genetic Variation. Genome Res. 9, 677–679.

53. Sudmant, P.H., Rausch, T., Gardner, E.J., Handsaker, R.E., Abyzov, A., Huddleston, J., Zhang, Y., Ye, K., Jun, G., Fritz, M.H.-Y., et al. (2015). An integrated map of structural variation in 2,504 human genomes. Nature 526, 75–81.

54. Taliun, D., Harris, D.N., Kessler, M.D., Carlson, J., Szpiech, Z.A., Torres, R., Taliun, S.A.G., Corvelo, A., Gogarten, S.M., Kang, H.M., et al. (2021). Sequencing of 53,831 diverse genomes from the NHLBI TOPMed Program. Nature 590, 290–299.

55. Telenti, A., Pierce, L.C.T., Biggs, W.H., di Iulio, J., Wong, E.H.M., Fabani, M.M., Kirkness, E.F., Moustafa, A., Shah, N., Xie, C., et al. (2016). Deep sequencing of 10,000 human genomes. Proc. Natl. Acad. Sci. U. S. A. 113, 11901–11906.

56. The 1000 Genomes Project Consortium (2010). A map of human genome variation from population-scale sequencing. Nature 467, 1061–1073.

57. The 1000 Genomes Project Consortium (2012). An integrated map of genetic variation from 1,092 human genomes. Nature 491, 56–65.

58. The 1000 Genomes Project Consortium (2015). A global reference for human genetic variation. Nature 526, 68–74.

59. The International HapMap 3 Consortium (2010). Integrating common and rare genetic variation in diverse human populations. Nature 467, 52.

60. Van der Auwera, G.A., and O’Connor, B.D. (2020). Genomics in the Cloud: Using Docker, GATK, and WDL in Terra (“O’Reilly Media, Inc.”).

61. Wagner, J., Olson, N.D., Harris, L., McDaniel, J., Khan, Z., Farek, J., Mahmoud, M., Stankovic, A., Kovacevic, V., Yoo, B., et al. (2021). Benchmarking challenging small variants with linked and long reads. bioRxiv 2020.07.24.212712.

62. Werling, D.M., Brand, H., An, J.-Y., Stone, M.R., Zhu, L., Glessner, J.T., Collins, R.L., Dong, S., Layer, R.M., Markenscoff-Papadimitriou, E., et al. (2018). An analytical framework for whole-genome sequence association studies and its implications for autism spectrum disorder. Nat. Genet. 50, 727–736.

63. Wigginton, J.E., Cutler, D.J., and Abecasis, G.R. (2005). A note on exact tests of Hardy-Weinberg equilibrium. Am. J. Hum. Genet. 76, 887–893.

64. Zhao, H., Sun, Z., Wang, J., Huang, H., Kocher, J.-P., and Wang, L. (2014). CrossMap: a versatile tool for coordinate conversion between genome assemblies. Bioinformatics 30, 1006–1007.

65. Zhao, X., Weber, A.M., and Mills, R.E. (2017). A recurrence-based approach for validating structural variation using long-read sequencing technology. Gigascience 6, 1–9.

66. Zheng-Bradley, X., and Flicek, P. (2017). Applications of the 1000 Genomes Project resources. Brief. Funct. Genomics 16, 163–170.

67. Zook, J.M., McDaniel, J., Olson, N.D., Wagner, J., Parikh, H., Heaton, H., Irvine, S.A., Trigg, L., Truty, R., McLean, C.Y., et al. (2019). An open resource for accurately benchmarking small variant and reference calls. Nat. Biotechnol. 37, 561–566.

